# Regulatory and coding sequences of TRNP1 co-evolve with brain size and cortical folding in mammals

**DOI:** 10.1101/2021.02.05.429919

**Authors:** Zane Kliesmete, Lucas E. Wange, Beate Vieth, Miriam Esgleas, Jessica Radmer, Matthias Hülsmann, Johanna Geuder, Daniel Richter, Mari Ohnuki, Magdalena Götz, Ines Hellmann, Wolfgang Enard

## Abstract

Brain size and cortical folding have increased and decreased recurrently during mammalian evolution. Identifying genetic elements whose sequence or functional properties co-evolve with these traits can provide unique information on evolutionary and developmental mechanisms. A good candidate for such a comparative approach is *TRNP1*, as it can control proliferation of neural progenitors in mice and ferrets. Here, we investigate the contribution of both regulatory and coding sequences of *TRNP1* to brain size and cortical folding in over 30 mammals. We find that the rate of TRNP1 protein evolution (*ω*) significantly correlates with brain size, slightly less with cortical folding and much less with body size. This brain correlation is stronger than for >95% of random control proteins. This co-evolution is likely affecting TRNP1 activity, as we find that TRNP1 from species with larger brains and more cortical folding induce higher proliferation rates in neural stem cells. Furthermore, we compare the activity of putative cis-regulatory elements (CREs) of *TRNP1* in a massively parallel reporter assay (MPRA) and identify one CRE that co-evolves with cortical folding in Old World Monkeys and Apes. Our analyses indicate that coding and regulatory changes that increased *TRNP1* activity were positively selected either as a cause or a consequence of increases in brain size and cortical folding. They also provide an example how phylogenetic approaches can inform biological mechanisms, especially when combined with molecular phenotypes across several species.

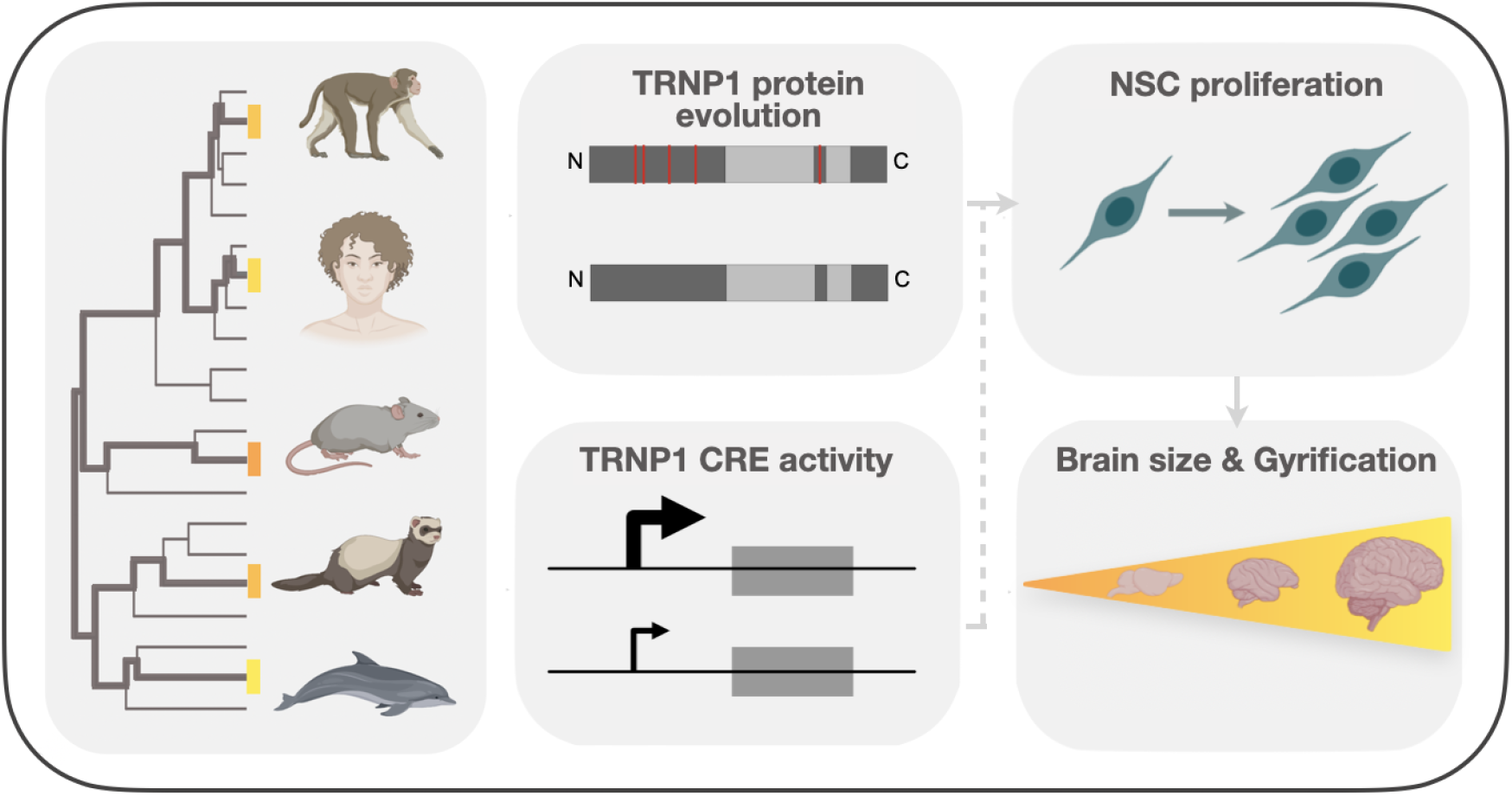

## Introduction

Understanding the genetic basis of complex phenotypes within and across species is central for biology. Brain phenotypes - even when as simple as size or folding-are of particular interest to many fields, because they are linked to cognitive abilities, which are of particular interest to humans^1,2^. Brain size and cortical folding show extensive variation across mammals, including recurrent independent increases and decreases^34,56^. For example, most rodents have a small brain and an unfolded cortex^7^, while carnivores, cetaceans and primates generally have enlarged and folded cortices, peaking in dolphin and human. Also within primates these traits vary, showing an increase on the great ape branch, but also decreases in several New World monkey species. Using comparative, i.e. phylogenetic, approaches across primates and mammals, these variations have been correlated with different life history traits, such as longevity, diet or energetic constraints^8,2,9^ revealing underlying ecological factors that drive selection for larger brains. The underlying genetic and cellular factors that are associated with these evolutionary variations in brain size and folding have not been studied across such large phylogenies. However, observational and experimental studies, especially in mice, but 14 increasingly also in other systems like the ferret, macaques and humans, have led to major insights into the genetic and cellular mechanisms of cortical development^10,11,12^. Briefly, proliferation of neuroepithelial stem cells (NECs) that have contacts with the apical surface and basal lamina, leads to the formation of the neuroepithelium during early development. NECs then become Pax6 positive apical progenitors (APs), i.e. apical Radial Glia Cells (aRGCs), that continue to self-amplify before producing basal progenitors (BPs). BPs include basal Radial Glia Cells (bRGCs) that remain Pax6 positive, loose the apical contact and - depending on the species - can also self-amplify before eventually producing neurons. The extent of proliferation in aRGCs and bRGCs is governed by their cell-cycle length where a short cell-cycle leads to symmetric divisions and delayed onset of neurogenesis, and these processes largely determine the cortical size and folding. Hence, genes that influence the proliferation of aRGC and/or bRGC are good candidates to contain genetic changes that contribute to evolutionary changes in brain size and folding. The major focus in this respect has been on identifying and functionally characterizing genetic changes on the human or primate lineage. For example, the human-specific paralog ARHGAP11B was found to induce bRGC proliferation and folding in cortices of mice, ferrets and marmosets^13,14,15^. Other examples include an amino-acid substitution specific to modern humans in *TKTL1*^16^, human-specific NOTCH2 paralogs^17,18^, the primate-specific genes TMEM14B and TBC1D3^19,20^ and an enhancer of *FZD8*, a receptor of the Wnt pathway^21^. While mechanistically convincing, it is unclear whether the proposed evolutionary link can be generalized as only one evolutionary lineage is investigated. While more evolutionary lineages are investigated in comparative approaches that correlate sequence changes with brain size changes^22,3^, these studies lack mechanistic evidence and are limited to the analysis of protein-coding regions. Here, we combine mechanistic and phylogenetic approaches to study *TRNP1*, a gene that is known to be important for cortical growth and folding by influencing aRGC and bRGC proliferation and differentiation in mice^23,24,25^ and ferrets^26^.

The presence of Trnp1 in neural stem cells (NSCs) isolated from mouse cortices induces phase separation, accelerating mitosis progression which in turn leads to increased self-renewal and proliferation and, ultimately, tangential expansion of the cortex^23^. For this especially important are the N-terminal intrinsically disordered regions^27^. Furthermore, knocking down Trnp1 in mice or expressing a dominant-negative TRNP1 in ferrets after the initial formation of aRGCs leads to more basal progenitors and subsequent folding of the cortex^23,24,26^. Recently, its expression in basal progenitors has also been linked to increased proliferation and cortical folding^25^. Altogether, there is strong mechanistic evidence that the structure and regulation of *TRNP1* are important for brain development in the investigated model organisms. However, it remains unclear what role *TRNP1* evolution played for brain evolution across a larger range of mammals. Here, we investigate the evolution of TRNP1 regulatory and coding sequences across mammals and infer to what extent these are linked to brain evolution.

## Results

### TRNP1 amino acid substitution rates co-evolve with brain size and cortical folding in mammals

We experimentally and computationally collected^28^ and aligned^29^ 45 mammalian TRNP1 coding sequences, including dolphin and 18 primates (99.0% completeness, Supplementary Figure 1A). Using this large multiple alignment, we find that the best fitting evolutionary model suggests that 9.8% of the codons show signs of recurrent positive selection (i.e. *ω* > 1, M8 vs. M7 model of PAML^30^, *χ*^2^ p-value< 0.001, df=2). Eight codons with a selection signature could be pinpointed with high confidence (Supplementary table 5). Seven out of those eight reside within the first intrinsically disordered region (IDR) and one in the second IDR of the protein (Figure 1B; Supplementary Figure 1B). The IDRs of TRNP1 are thought to mediate homotypic and heterotypic protein-protein interactions and are relevant for TRNP1-dependent phase separation, nuclear compartment size regulation and M-phase length regulation^27^. Hence, the positively selected sites indicate that these IDR-mediated TRNP1 functions were repeatedly adapted during mammalian evolution and the identified sites are candidates for further functional studies. However, even though this analysis shows that TRNP1 evolved under positive selection, it is yet unclear why.

**Figure 1.**
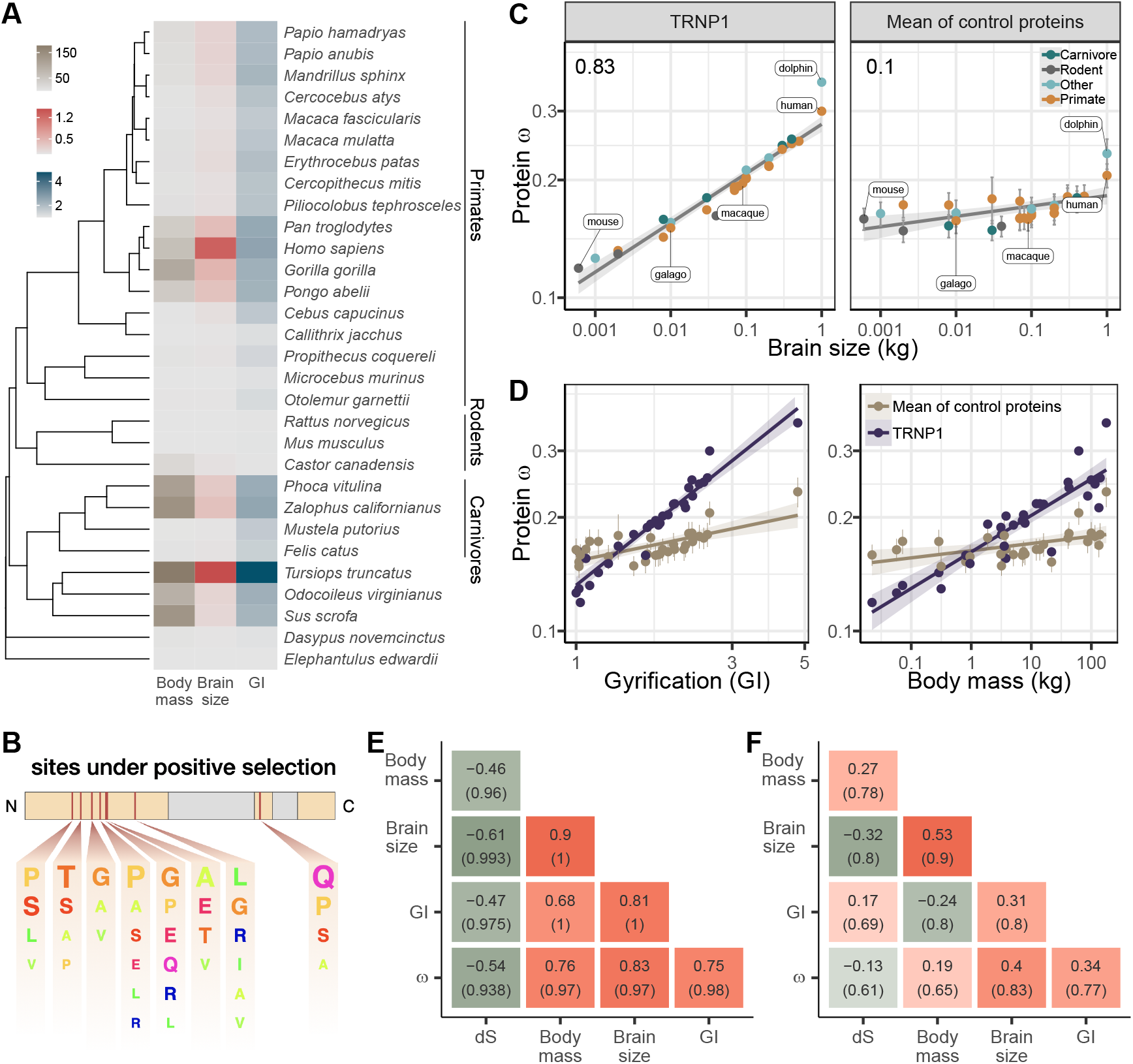
TRNP1 amino acid substitution rates co-evolve with brain size and cortical folding in mammals. (*A*) Mammalian species for which body mass, brain size, GI measurements and TRNP1 coding sequences were available (n=30). Units: body mass and brain size in kg; GI is a ratio (cortical surface / perimeter of the brain surface). (*B*) Scheme of the mouse TRNP1 protein (223 AAs) with intrinsically disordered regions (orange) and sites (red lines) subject to positive selection in mammals (*ω* > 1, *pp* > 0.95; Supplementary Figure 1B). The letter size of the depicted AAs represents the abundance of AAs at the positively selected sites. (*C*) TRNP1 protein substitution rates (*ω*) significantly correlate with brain size (*r* = 0.83, *pp*=0.97, left), which is considerably higher than the average correlation across 125 control proteins (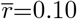, right). (D) TRNP1 *ω* correlation with GI (left) and body mass (right) compared to the average across control proteins. (*C, D*) Errorbars indicate standard errors. (*E, F*) Estimated marginal (*E*) and partial (*F*) correlation matrices of the combined model including the three traits and substitution rates. Posterior probabilities of the associations are depicted in brackets.

One approach to tackle this question is to correlate the rate of TRNP1 protein evolution (ω) with the rate of cortical folding and brain size evolution. To this end we compiled phenotypic data on brain size, cortical folding and body mass that was available for 30 mammalian species (Figure 1A; Supplementary table 4). Here, brain size is quantified as its weight and cortical folding as the ratio of the cortical surface over the perimeter of the brain surface, the gyrification index (GI), where a GI= 1 indicates a completely smooth brain and a GI> 1 indicates higher levels of cortical folding^31^.

The matter is complicated by the high correlation among the brain phenotypes that are of interest - brain size and GI - and body mass as it is unclear whether this is caused by co-varying selection pressures and/or developmental constraints^32,6^. In addition, the rate of protein evolution (*ω*) also varies due to the efficacy of selection, which in turn varies across the branches of the phylogeny due to differences in the effective population size^33,34^, which varies with body size^35,36^. In order to control for this, we compiled a set of 125 control proteins with similar length, number of exons and alignment quality across the 30 species (Supplementary Figure 2, Methods) to estimate the average correlation between brain size/GI and *ω*. We also included body mass to further control for the specificity of our findings^35,36^. Thus, if brain size and GI show a stronger correlation with *ω* of TRNP1 than the average association across the control proteins and also a stronger correlation than with body mass, this would suggest that the link between the brain traits and protein evolution is at least in part due to direct selection.

In order to test whether the evolution of the TRNP1 protein coding sequences is linked to any of the three traits, we used Coevol^36^, a Bayesian MCMC method that jointly models the rates of substitutions and quantitative traits. The resulting covariance matrix of substitution rates (branch length *λ_S_*, *ω*) and the phenotypic traits then allows for a quantitative evaluation of a potential co-evolution using the posterior probability (*pp*) of the correlations^36^. Considering the traits separately, we find that brain size has the highest marginal correlation with *ω* (*r*=0.789, *pp*=0.97), followed by GI (*r*=0.769, *pp*=0.98), and body mass (*r*=0.696, *pp*=0.95) (Supplementary table 6). To better disentangle their effects, we then simultaneously inferred their correlations (Figure 1C-F; Supplementary table 6). Brain size remained the strongest marginal correlation (*r*=0.827, *pp*=0.97) and also the strongest partial correlation (*r*=0.403, *pp*=0.83) which was slightly higher than that with GI (*r*=0.341, *pp*=0.77) and considerably stronger than the one with body mass (*r*=0.19, *pp*=0.65). Moreover, the observed association with brain size is much higher than the average association across the control protein sequences (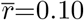, Figure 1C right), validating that this is not caused by the effective population size differences across the species. TRNP1 partial correlation with brain size is within the highest 4% observed across the 126 proteins and correlation with GI among the highest 6.3%, while the association with body size is only among the top 13.5%, again supporting a rather direct link between TRNP1 and brain phenotypes (Supplementary Figure 2). In addition, only 3 other proteins (<3%) with brain phenotype association showed higher a significant proportion of sites evolving under positive selection across the phylogeny (Supplementary Figure 3). Hence, these results show that TRNP1 evolved under positive selection and that its rate of sequence evolution is linked to the evolution of brain size and cortical folding.

### TRNP1 proliferative activity coevolves with brain size and cortical folding in mammals

Next, we investigated whether the correlation between TRNP1 protein evolution and cortical phenotypes can be linked to functional properties of TRNP1 at a cellular level. A central property of TRNP1 is to promote proliferation of aRGC^23,27^ and also of BPs^25^. This proliferative activity can be assessed in an *in vitro* assay in which *Trnp1* is transfected into neural stem cells (NSCs) isolated from E14 mouse cortices^23,27^. To compare TRNP1 orthologoues in this assay, we synthesized and cloned the Trnp1 coding sequence of human, rhesus macaque, galago, mouse and dolphin that cover the observed range of *ω* (Figure 1C). After co-transfection with GFP, we quantified the number of proliferating (Ki67+, GFP+) over all transfected (GFP+) NSCs for each *TRNP1* orthologue in ≥ 7 replicates (Figure 2A,B). We confirmed that *TRNP1* transfection does increase proliferation compared to a GFP-only control (*p*-value< 2 × 10^-16^; Supplementary Figure 4A) as shown in previous studies^23,27^. Remarkably, the proportion of proliferating cells was highest in cells transfected with dolphin TRNP1 followed by human, which was significantly higher than the two other primates, galago and macaque (Figure 2C; Supplementary Figure 4B; Supplementary tables 7–9). Indeed, the proliferative activity of TRNP1 is a significant predictor for brain size (*p*-value= 0.001, *R*^2^ = 0.89) and GI (*p*-value= 0.016, *R*^2^ = 0.69) in its species of origin (Phylogenetic generalised least squares PGLS, Likelihood Ratio Test (LRT); Figure 2C). Note that the three primates and the dolphin are phylogenetically equally distant to the mouse (Figure 2C) and hence a bias due to the murine assay system can not explain the observed correlations with brain size and GI. Hence, these results further support that the TRNP1 protein co-evolves with brain size and cortical folding.

**Figure 2.**
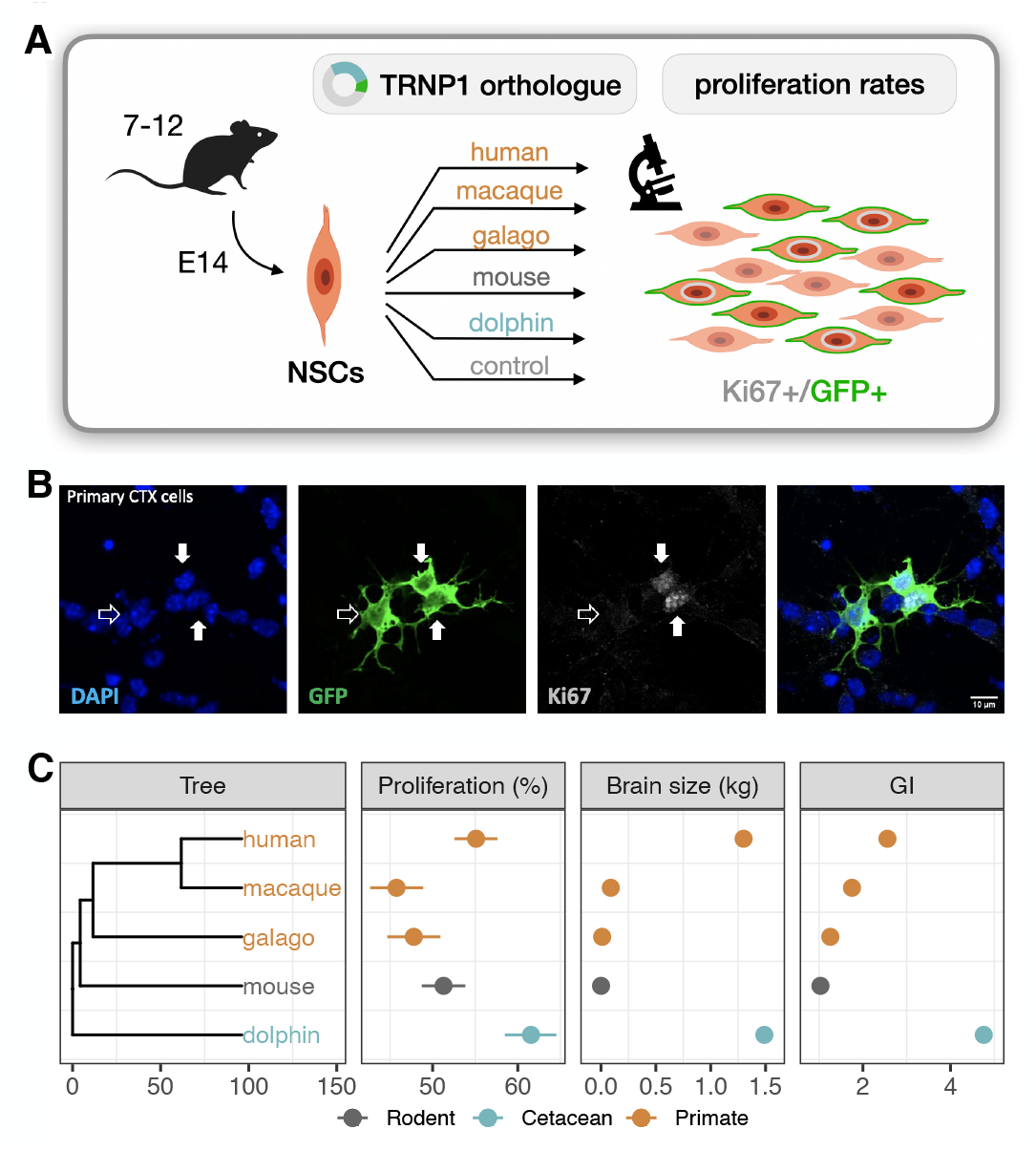
TRNP1 proliferative activity correlates with brain size and cortical folding. (*A*) Five different TRNP1 orthologues were transfected into neural stem cells (NSCs) isolated from cerebral cortices of 14 day old mouse embryos and proliferation rates were assessed after 48 h using Ki67-immunostaining as proliferation marker and GFP as transfection marker in 7-12 independent biological replicates. (*B*) Representative image of the transfected cortical NSCs immunostained for GFP and Ki67. Arrows indicate three transfected cells of which two (solid arrows) are Ki67-positive. (*C*) Induced proliferation in NSCs transfected with TRNP1 orthologues from 5 different species (Table 8). Proliferation rates are a significant predictor for brain size (*χ*^2^=10.04, df=1, *p*-value= 0.001; *β*=11.75 ±2.412, *R*^2^ = 0.89) and GI (*χ*^2^=5.85, df=1, *p*-value= 0.016; *β*=16.97 ± 6.568, *R*^2^ = 0.69) in the respective species (PGLS, LRT). Error bars indicate standard errors.

### Activity of a cis-regulatory element of *TRNP1* co-evolves with cortical folding in catarrhines

Experimental manipulation of Trnp1 expression levels alters proliferation and differentiation of aRGC and bRGC in mice and ferrets^23,26,25^. Hence, we next investigated whether changes in *TRNP1* regulation may also be associated with the evolution of cortical folding and brain size by analyzing co-variation in the activity of *TRNP1* associated cis-regulatory elements (CREs), using a massively parallel reporter assays (MPRAs). To this end, a library of putative regulatory sequences is cloned into a reporter vector and their activity is quantified simultaneously by the expression levels of element-specific barcodes^37^. To identify putative CREs of *TRNP1*, we used DNase Hypersensitive Sites (DHS) from human fetal brain^38^ and found three upstream CREs, the promoter-including exon 1, an intron CRE, one CRE overlapping the second exon, and one downstream CRE (Figure 3A). We obtained the orthologous sequences of the human CREs using reciprocal best blat strategy across additional mammalian species either from genome databases or by sequencing, yielding a total of 351 putative CREs in a panel of 75 mammalian species (Supplementary Figure 5). Due to limitations in the length of oligonucleotide synthesis, we cut each orthologous putative CRE into highly overlapping, 94 bp fragments. The resulting 4950 sequence tiles were synthesised together with a barcode unique for each tile. From those, we constructed a complex and unbiased lentiviral plasmid library containing at least 4251 (86%) CRE sequence tiles (Figure 3B,C). Next, we stably transduced this library into neural progenitor cells (NPCs) derived from two humans and one macaque^39^. We calculated the activity per CRE sequence tile as the read-normalised reporter gene expression over the read-normalised input plasmid DNA (Figure 3A, Methods). Finally, we use the per-tile activities (Supplementary Figure 6A) to reconstruct the activities of the putative CREs. To this end, we summed all tile sequence activities for a given CRE while correcting for the built-in sequence overlap (Figure 3D; Methods). CRE activities correlate well within the two human NPC lines and between the human and macaque NPC lines, indicating that the assay is robust across replicates and species (Pearson’s *r* 0.85-0.88; Supplementary Figure 6B). The CREs covering exon 1, the intron and the CRE downstream of *TRNP1* show the highest total activity across species while the CREs upstream of *TRNP1* show the lowest activity (Figure 3E).

**Figure 3.**
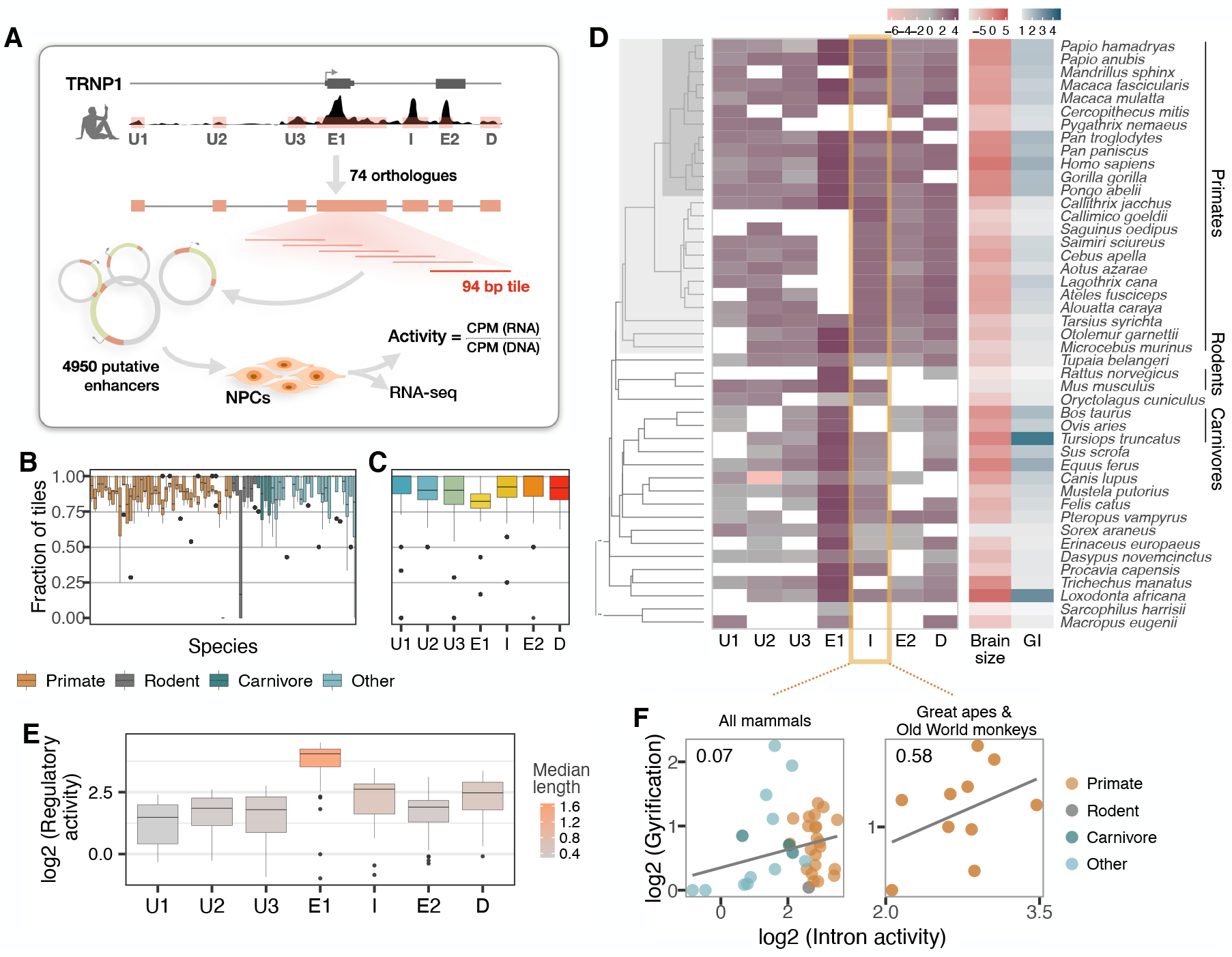
Activity of a cis-regulatory element (CRE) of *TRNP1* correlates with cortical folding in catarrhines. (*A*) Experimental setup of the MPRA assay. Regulatory activity of 7 putative TRNP1 CREs from 75 species were assayed in neural progenitor cells (NPC) derived from human and macaque induced pluripotent stem cells. (*B*) Fraction of the detected CRE tiles in the plasmid library per species across regions. The detection rates are unbiased and uniformly distributed across species and clades with only one extreme outlier *Dipodomys ordii*. (*C*) Fraction of the detected CRE tiles in the plasmid library per region across species. (*D*) Log-transformed total regulatory activity per CRE in human NPCs across species with available brain size and GI measurements (n=45). (*E*) Total activity per CRE across species. Exon1 (E1), intron (I) and the downstream (*D*) regions are more active and longer than other regions. (*B,C,E*) Each box represents the median and first and third quartiles with the whiskers indicating the furthest value no further than 1.5 * IQR from the box. Individual points indicate outliers. (*F*) Regulatory activity of the intron CRE is moderately associated with gyrification across mammals (PGLS, LRT *p*-value< 0.1, *R*^2^ = 0.07, n=37) and strongest across great apes and Old World Monkeys, i.e. catarrhines (PGLS, LRT *p*-value< 0.003, R^2^ = 0.58, n=10).

Next, we tested whether CRE activity is associated with either brain size or GI across the 45 of the 75 mammalian species for which these phenotypes were available (Figure 3D). None of the CREs showed any association with brain size (PGLS, LRT p-value> 0.1). In contrast, we found that the CRE activity of the intron CRE had a slight positive association with gyrification (PGLS, LRT *p*-value< 0.1, Figure 3F left; Table 10). These associations are much weaker than those observed with TRNP1 protein evolution. Part of the reason might be that CREs have a much higher evolutionary turnover rate than coding sequences^40,41,42^, which would result in orthologous DNA sequences that do not function as CREs or even to a loss of the orthologous sequences. The latter effect might explain why the sequences orthologous to human CREs are shorter in species more distantly related to humans (Supplementary Figure 5).

Therefore, we restricted our analysis to the catarrhine clade that encompasses Old World Monkeys, great apes and humans. Here, the association between intron CRE activity and GI becomes considerably stronger (PGLS, LRT *p*-value< 0.003, Figure 3F right; Table 11). Moreover, the intron CRE activity-GI association was consistently detected across all three cell lines including the macaque NPCs (Table 11), which is consistent with its identification as an enhancer in developing human and macaque cortices^43^. Hence, the observed correlation is unlikely to be caused by a bias in the assay, indicating that also the expression regulation of Trnp1 co-evolves with cortical folding at least in catarrhines.

### Transcription factors with binding site enrichment on intron CREs regulate cell proliferation and explain the observed activity across catarrhines

Reasoning that differences in CRE activities will likely be mediated by differences in their interactions with transcription factors (TF), we analysed the sequence evolution of putative TF binding sites (Figure 4A). First, we performed RNA-seq on the same samples that were used for the MPRA assay. Notably, also *TRNP1* was expressed (Figure 4B), supporting the relevance of our cellular system. Among the 392 expressed TFs with known binding motifs, we identified 22 with an excess of binding sites^44^ within the catarrhine intron CRE sequences (Figure 4B,D). In agreement with TRNP1 itself being involved in the regulation of cell proliferation^45,23,27^, these 22 TFs are enriched in biological processes regulating cell proliferation, neuron apoptotic process and hormone levels (Figure 4C; Table 12).

**Figure 4.**
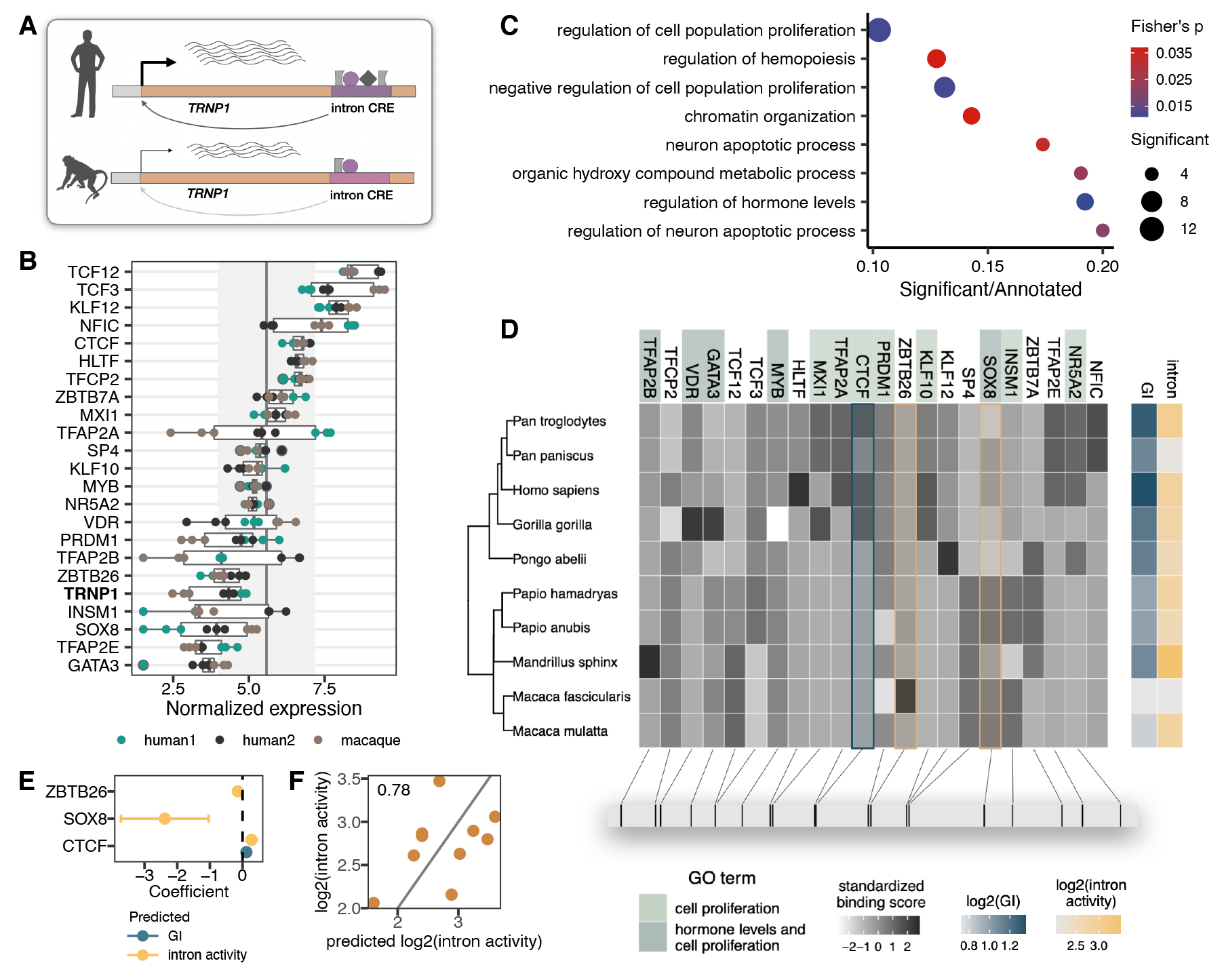
Transcription factors with binding site enrichment on intron CREs regulate cell proliferation and explain the observed activity across catarrhines. (*A*) Orthologous intron CRE sequences show different regulatory activities under the same cellular conditions, suggesting variation in cis-regulation across species. (*B*) Variance-stabilized expression in NPCs of *TRNP1* and the 22 transcription factors (TFs) with enriched binding sites (motif weight>=1) on the intron CREs. Each box represents the median, first and third quartiles with the whiskers indicating the furthest value no further than 1.5 * IQR from the box. Points indicate individual expression values. Vertical line indicates average expression across all 392 TFs (5.58), grey area: standard deviation (1.61). (*C*) Eight top enriched biological processes (Gene Ontology, Fisher’s *p*-value< 0.05) of the 22 TFs. Background: all expressed TFs (392). (*D*) Variation in binding scores of the enriched TFs across catarrhines. Heatmaps indicate standardised binding scores (grey), GI values (blue) and intron CRE activities (yellow) from the respective species. TF background color indicates gene ontology assignment of the TFs to the 2 most significant biological processes. The bottom panel indicates the spatial position of the top binding site (motif score>3) for each TF on the human sequence. (*E*) Binding scores of 3 TFs (CTCF, ZBTB26, SOX8) are predictive for intron CRE activity, whereas only CTCF binding shows an association with the GI (PGLS, LRT *p*-value< 0.05). (*F*) Predicted intron CRE activity by the binding scores of the 3 TFs vs the measured intron CRE activity across catarrhines.

To further prioritise these 22 TFs, we used the motif binding scores in the 10 catarrhine intron CREs to predict the observed intron CRE activity in the MPRA assay and to predict the GI of the respective species. We found three TFs (CTCF, ZBTB26, SOX8) to be predictive for intron CRE activity and one TF (CTCF) to be predictive for GI (PGLS, LRT *p*-value< 0.05, Figure 4D-F). In summary, we find evidence that a higher activity of the intron CRE is correlated with gyrification in catarrhines, indicating that also regulatory changes of *TRNP1* contributed to the evolution of gyrification.

## Discussion

Previous studies in mice and ferrets have elucidated mechanisms how *Trnp1* is necessary for proliferation and differentiation of neural progenitors and how it could contribute to the evolution of brain size and cortical folding. We applied phylogenetic methods to find associations between sequence and trait evolution and found that the rate of protein evolution and the proliferative activity of TRNP1 positively correlate with brain size and gyrification in mammals and that the activity of a regulatory element in the intron of *TRNP1* positively correlates with gyrification in catarrhines. At the sequence level, such a correlation could also be caused by confounding factors that affects the efficacy of natural selection such as the effective population size^33,34^. However, body size - a reasonable proxy for effective population size^35,36^-, correlates much less with TRNP1 protein evolution than brain size or gyrification. Even more convincingly, the correlation of TRNP1 with brain size and GI is much stronger than the correlation of these traits with the evolution of other proteins, that would have had to experience the same population size changes. Furthermore, it is unclear how an increased proliferative activity of TRNP1 or an increased CRE activity could be caused by a reduced efficacy of selection or other confounding factors. Together with the known role of TRNP1 in brain development, we think that the observed correlations are best interpreted as co-evolution of TRNP1 activity with brain size and gyrification, i.e. that more active TRNP1 alleles were selected because they were advantageous to increase brain size and/or gyrification.

Of note, the effect of structural changes appears stronger than the effect of regulatory changes. This is contrary to the notion that regulatory changes should be the more likely targets of selection as they are more cell-type specific^46^ (but see also^47^). However, current measures of regulatory activity are inherently less precise than counting amino acid changes, which will necessarily deflate the estimated association strength^40,41,42^. In any case, our analysis suggests that evolution combined both regulatory and structural evolution to modulate TRNP1 activity.

The MPRA also allowed to identify transcription factors that evolved a binding site enrichment to the intron CRE and are likely direct regulators of TRNP1. These include INSM1^48^, which also has been shown to control NSC-to-neural-progenitor transition, as well as other relevant factors with increased activity in human neural stem and progenitor cells during early cortical development compared to later stages, such as TFAP2AIC, TCF3, KLF12 and again INSM1^49,50^. Among the enriched TFs that bind to the intron CRE, CTCF had the strongest association with gyrification. Although CTCF is best known for its insulating properties, it can also act as transcriptional activator and recruit co-factors in a lineage-specific manner^51^.In NSCs, CTCF loss causes severe impairment in proliferative capacity through the increase in premature cell-cycle exit, which results in drastically reduced NSC pool and early differentiation [52]. The overlapping molecular roles of TRNP1 and CTCF in NSCs support the possibility that TRNP1 is among the cell-fate determinants downstream of CTCF^53,54^. Differences between species in CTCF binding strength and/or length to the intron CRE might have direct consequences for the binding of additional TFs, TRNP1 expression and the resulting NSC pool. However, the effects of CTCF binding *in vitro* and *in vivo* might differ and the exact mechanism, including the developmental timing and cellular context in which this is relevant, is yet to be disentangled.

Independent from the mechanisms and independent whether caused by regulatory or structural changes, it is relevant how an increased TRNP1 activity could alter brain development. When overexpressing *Trnp1* in aRGCs of developing mice (E13) and ferrets (E30), aRGC proliferation increases^23,24,26^. Similarly, overexpression of *Trnp1* increases proliferation *in vitro* in NSCs^23,27^ or breast cancer cells^45^. Hence, Trnp1 evolution could contribute to evolving a larger brain by increasing the pool of aRGCs. In addition, increases in brain size and especially increases in cortical folding are highly dependent on increases in proliferation of basal progenitors (BPs), in particular bRGCs^10,11,12^. Remarkably, recent evidence indicates that Trnp1 could be important also for the proliferation of BPs^25^: Firstly, in contrast to non-proliferating BPs from mice, proliferating BPs from human do express TRNP1^25^. Furthermore, when activating expression of *Trnp1* using CRISPRa at E14.5, more proliferating BPs and induction of cortical folding is observed^25^. Hence, a more active TRNP1 can increase proliferation in aRGCs and BPs and this could cause the observed co-evolution with brain size and cortical folding. *TRNP1* is the first case where analyses of protein sequence, regulatory activity and protein activity across a larger phylogeny have been combined to investigate the role of a candidate gene in brain evolution. Functional evidence from evolutionary changes on the human lineage e.g. for ARHGAP11B and NOTCH2NL. but also phylogenetic evidence from correlating sequence changes with brain size changes^3,22^ indicate that a substantial number of genes could adapt their function when brain size changes in mammalian lineages. Improved genome assemblies^55^ will decisively improve phylogenetic approaches^56,57,58,59^. In combination with the increased possibilities for functional assays due to DNA synthesis^60^ and comparative cellular resources across many species^61,62,39^ this offers exciting possibilities to study the genetic basis of complex phenotypes within and across species.

## Material and Methods

### Sample collection and cell culture

#### Primary cerebral cortex harvesting and culture

E14 Mouse (Mus musculus) cerebral cortices were dissected, removing the ganglionic eminence, the olfactory bulb, the hippocampal anlage and the meninges. Cells were mechanically dissociated with a fire polish Pasteur pipette. Cells were then seeded onto poly-D-lysine (PDL)-coated glass coverslips in DMEM-GlutaMAX (Dulbeccos’s modified Eagles’s medium) supplemented with 10% fetal calf serum (FCS) and 100 μg/ml penicilin-streptomycin (Pen.Strep.) and cultured at 37°C in a 5% CO^2^incubator.

#### Culture of HEK293T cells

HEK 293T cells (Homo sapiens) were grown in DMEM supplemented with 10% FCS and 1% Pen. Strep. Cells were cultured in 10 cm flat-bottom dishes at 37°C in a 5% CO^2^ environment and split every two to three days in a 1:10 ratio using 5 mL PBS to wash and 0.5 mL 0.25 % Trypsin to detach the cells.

#### Culture of Neuro-2A cells

Neuro-2A cells (N2A) (ATCC; CCL-131, Mus musculus) were culutred in Eagle’s Minimum Essential Medium (EMEM, Thermo Fischer Scientific) with 10% FCS (Thermo Fischer Scientific) at 37°C in a 5% CO^2^ incubator and split every two to three days in a 1:5 ratio using 5 mL PBS (Thermo Fischer Scientific) to wash and 0.5 mL 0.25 % Trypsin (Thermo Fischer Scientific) to detach the cells.

#### Culture of neural progenitor cells

Neural progenitor cells of two human (Homo sapiens) and one cynomologous monkey (Macaca fascicularis) cell line [39], were cultured at 37°C in a 5% CO^2^ incubator on Geltrex (Thermo Fisher Scientific) in DMEM F12 (Fisher scientific) supplemented with 2 mM GlutaMax-I (Fisher Scientific), 20 ng/μl bFGF (Peprotech), 20 ng/μl hEGF (Miltenyi Biotec), 2% B-27 Supplement (50X) minus Vitamin A (Gibco), 1% N2 Supplement 100X (Gibco), 200*μ*M L-Ascorbic acid 2-phosphate (Sigma) and 100 *μ*g/ml penicillin-streptomycin with medium change every second day. For passaging, NPCs were washed with PBS and then incubated with TrypLE Select (Thermo Fisher Scientific) for 5 min at 37°C. Culture medium was added and cells were centrifuged at 200 × g for 5 min. Supernatant was replaced by fresh culture medium and cells were transferred to a new Geltrex coated dish. The cells were split every two to three days in a ratio of 1:3.

### Sequencing of *TRNP1* for primate species

#### Identification of cis-regulatory elements of *TRNP1*

DNase hypersensitive sites in the proximity to *TRNP1* (25 kb upstream, 3 kb downstream) were identified in human fetal brain and mouse embryonic brain DNase-seq data sets^63,38^ downloaded from NCBI’s Sequence Read Archive (See Key Resources Table). Reads were mapped to human genome version hg19 and mouse genome version mm10 using NextGenMap with default parameters (NGM; v. 0.0.1)^64^. Peaks were identified with Hotspot v.4.0.0 using default parameters^65^. Overlapping peaks were merged, and the union per species was taken as putative cis-regulatory elements (CREs) of *TRNP1* (Table 2). The orthologous regions of human *TRNP1* DNase peaks in 49 mammalian species were identified with reciprocal best hit using BLAT (v. 35×1)^66^. Firstly, sequences of human *TRNP1* DNase peaks were extended by 50 bases down and upstream of the peak and the best matching sequence per peak region were identified with BLAT using the following settings: -t=DNA -q=DNA -stepSize=5 -repMatch=2253 -minScore=0 -minIdentity=0 -extendThroughN. These sequences were aligned back to hg19 using the same settings as above. The resulting best matching hits were considered reciprocal best hits if they fell into the original human *TRNP1* CREs. In total 351 putative TRNP1 CRE sequences were identified, including human, mouse and ortholous sequences.

#### Cross-species primer design for sequencing

We sequenced TRNP1 coding sequences in 6 primates for which reference genome assemblies were either unavailable or very sparse and the ferret (*Mustela putorius furo*) where the sequence was incomplete (see Table 1). For the missing primate sequences we used NCBI’s tool Primer Blast^67^ with the human *TRNP1* gene locus as a reference. Primer specificity was confirmed using the predicted templates in 12 other primate species available in Primer Blast. Following primers were used as they worked reliably in all 6 species (Forward primer, GGGAGGAGTAAACACGAGCC; Reverse Primer, AGCCAGGTCATTCACAGTGG). For the ferret sequence, the genome sequence (MusPutFur1.0,) contained a gap in the TRNP1 coding sequence leading to a truncated protein. To recover the full sequence of TRNP1 we used the conserved sequence 5’ of the gap and 3’ of the gap as input for primer blast (Primer sequences can be found in the analysis github, See Data and Code Availability).

In order to obtain *TRNP1* CREs for the other primate species, we designed primers using primux^68^ based on the species with the best genome assemblies and subsequently tested them in closely related species in multiplexed PCR reactions. A detailed list of designed primer pairs per CRE and reference genome can be found in the analysis github (see Data and Code Availability)

#### Sequencing of target regions for primate species

Primate gDNAs were obtained from Deutsches Primaten Zentrum, DKFZ and MPI Leipzig (see Table 3). Depending on concentration, gDNAs were whole genome amplified prior to sequencing library preparation using GenomiPhi V2 Amplification Kit (Sigma). After amplification, gDNAs were cleaned up using SPRI beads (CleaNA). Both *TRNP1* coding regions and CREs were resequenced starting with a touchdown PCR to amplify the target region followed by a ligation and Nextera XT library construction. *TRNP1* coding regions were sequenced as 250 bases paired end with dual indexing on an Illumina MiSeq, the CRE libraries libraries were sequenced 50bp paired end on an Illumina Hiseq 1500.

#### Assembly of sequenced regions

Reads were demultiplexed using deML^69^. The resulting sequences per species were subsequently trimmed to remove PCR-handles using cutadapt (version 1.6)^70^. For sequence reconstruction, Trinity (version 2.0.6) in reference-guided mode was used^71^. The reference here is defined as the mapping of sequences to the closest reference genome with NGM (version 0.0.1)^64^. Furthermore, read normalisation was enabled and a minimal contig length of 500 was set. The sequence identity of the assembled contigs was validated by BLAT^66^ alignment to the closest reference *TRNP1* as well as to the human *TRNP1.* The assembled sequence with the highest similarity and expected length was selected per species.

The same strategy was applied to the resequenced ferret genomic sequence, except that we used bwa-mem2^72^ for mapping and for the assembly with Trinity we set minimal contig length to 300 (reference genome musFur1). Only the part covering the 3’ end (specifically, the last 107 AAs) was successfully assembled, however, luckily, MusFur1 genome assembly already provides a good-quality assembly for the 5’ end of the protein. The overlapping 36 AAs (108 nucleotides) between both sources had a 100% agreement on the nucleotide sequence level, hence we collapsed the sequences from both sources to yield a full-length protein-coding sequence. In a neighbor joining tree, where we included the nucleotide sequences from all 30 mammalian TRNP1 orthologues, ferret sequence was placed within the other Carnivore sequences (between cat and a branch leading to seal, sea lion) as expected given the pylogenetic relationships of these species.

#### *TRNP1* coding sequence retrieval and alignment

Human TRNP1 protein sequence was retrieved from UniProt database^73^ under accession number Q6NT89. We used the human TRNP1 in a tblastn^28^ search of genomes from 45 species, without any repeat masking specified in Table 1 (R-package rBLAST version 0.99.2). The resulting sequences were re-aligned with PRANK^29^ (version 150803), using the mammalian tree from Bininda-Emonds et al.^74^.

#### Control gene set selection and alignment

Control genes were selected using consensus coding-sequence (CCDS) dataset for human GRCh38.p12 genome (35,138 coding-sequences, release 23)^75^. Reciprocal best blat^66^ (RBB) strategy was applied to identify the orthologous sequences in the other 29 species using -q=prot -t=dnax blat settings. We picked the best matching sequence per CDS in each species using a score based on the BLOSUM62 substitution matrix^76^ and gapOpening=3, gapExtension=1 penalties, and requiring at least 30% of the human sequence to be found in the other species. This sequence was extracted and the same strategy was applied when blatting the orthologous sequence to the human genome. If the target sequence with the best score overlaps at least 10% of the original CDS sequence positions, it was kept. To have a comparable gene set to TRNP1 in terms of statistical power and alignment quality, we selected all genes that had a similar human coding-sequence length as TRNP1 (<1000 nucleotides) and 1 coding exon (322). If RBB returned multiple matches per species per sequence with the same highest alignment score to the human sequence, we kept these only if the matching sequences were identical, which resulted in 274 genes. We further filtered for genes with all orthologous sequences of length at least 50% and below 200% relative to the length of the respective human protein-coding orthologue (257 genes). These were aligned using PRANK^29^ as for TRNP1, and manually inspected. 112 alignments were optimal, and we could get additional 22 high-quality alignments by searching orthologues in additional genome versions using the previously described RBB stategy (gorilla gorGor5.fa, dolphin GCF_011762595.1_mTurTru1, wild boar GCF_000003025.6_Sscrofa11.1, rhesus macaque GCF_003339765.1_Mmul_10, olive baboon GCA_000264685.2_Panu_3.0) and redoing the alignment.

### Evolutionary sequence analysis

#### Identification of sites under positive selection

Program codeml from PAML soft-ware^30^ (version 4.8) was used to infer whether a significant proportion of TRNP1 protein sites evolve under positive selection across the phylogeny of 45 species. Site models M8 and M7 were compared^77^, that allow *ω* to vary among sites across the phylogenetic tree, but not between branches. M7 and M8 are nested with M8 allowing for sites under positive selection with *ω_s_.* Likelihood ratio test (LRT) was used to compare these models. Naive Empirical Bayes (NEB) analysis was used to identify the specific sites under positive selection (Pr(*ω* > 1) > 0.95). The same analysis was also applied to the 125 control proteins and TRNP1 across 30 species to compare the signals of positive selection.

#### Inferring correlated evolution using Coevol

Coevol^36^ (version 1.4) was utilised to infer the covariance between TRNP1 and cotrol protein evolutionary rate ω with three morphological traits (brain size, GI and body mass) across species (Table 4). For each model, the MCMC was run three times for at least 10,000 cycles, using the first 1,000 as burn-in. For TRNP1 and 125 control proteins all parameters have a relative difference < 0.3 and effective size > 50, indicating good convergence, 9 control proteins did not reach such a convergence and were thereby excluded from further analyses. We report the average posterior probabilities (*pp*), the average marginal and partial correlations of the full model (Table 6) and the separate models where including only either one of the three traits (Table 6). The posterior probabilities for a negative correlation are given by 1 – *pp*. These were back-calculated to make them directly comparable, independently of the correlation direction, i.e. higher *pp* means more statistical support for the respective correlation.

### Proliferation assay

#### Plasmid construction

The five *TRNP1* orthologous sequences containing the restriction sites BamHI and XhoI were synthetized by GeneScript. All plasmids for expression were first cloned into a pENTR1a gateway plasmid described in Stahl et al., 2013^23^ and then into a Gateway (Invitrogen) form of pCAG-GFP (kind gift of Paolo Malatesta). The gateway LR-reaction system was used to then sub-clone the different TRNP1 orthologues into the pCAG destination vectors.

#### Primary cerebral cortex transfection

Primary cerebral cortex cultures were established as outlined under Experimental Model and Subject Details. Plasmids were transfected with Lipofectamine 2000 (Life technologies) according to manufacturer’s instruction 2h after seeding the cells onto PDL coated coverslips. One day later cells were washed with phosphate buffered saline (PBS) and then fixed in 4% Paraformaldehyde (PFA) in PBS and processed for immunostaining.

#### Immunostaining

Cells plated on poly-D-lysine coated glass coverslips were blocked with 2% BSA, 0.5% Triton-X (in PBS) for 1 hour prior to immunostaining. Primary antibodies (chicken alpha-GFP, Aves Labs: GFP-1010 and rabbit *alpha*-Ki67, abcam: ab92742) were applied in blocking solution overnight at 4°C. Fluorescent secondary antibodies were applied in blocking solution for 1 hour at room temperature. DAPI (4’.6-Diamidin-2-phenylindol, Sigma) was used to visualize nuclei. Stained cells were mounted in Aqua Polymount (Polysciences). All secondary antibodies were purchased from Life Technologies. Representative high quality images were taken using an Olympus FV1000 confocal laser-scanning microscope using 20×/0.85 N.A. water immersion objective. Images used for quantification were taken using an epifluorescence microscope (Zeiss, Axio ImagerM2) equipped with a 20×/0.8 N.A and 63×/1.25 N.A. oil immersion objectives. Post image processing with regard to brightness and contrast was carried out where appropriate to improve visualization, in a pairwise manner.

#### Proliferation rate calculation using logistic regression

The proportion of successfully transfected cells that proliferate under each condition (Ki67-positive/GFP-positive) was modeled using logistic regression (R-package stats (version 4.0.3), `glm function`) with logit link function 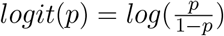, for 0≤p≤1, where *p* is the probability of success. The absolute number of GFP-positive cells were added as weights. Model selection was done using LRT within anova function from stats. Adding the donor mouse as a batch improved the models (Tables 7).

To back-calculate the absolute proliferation probability (i.e., rate) under each condition, intercept of the respective model was set to zero and the inverse logit function 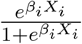 was used, where *i* indicates condition (Table 8). Two-sided multiple comparisons of means between the conditions of interest were performed using glht function (Tukey test, user-defined contrasts) from R package `multcomp` (version 1.4-13) (Table 9).

#### Phylogenetic modeling of proliferation rates using generalized least squares (PGLS)

The association between the induced proliferation rates for each TRNP1 orthologue and the brain size or GI of the respective species was analysed using generalised least squares (R-package `nlme`, version 3.1-143), while correcting for the expected correlation structure due to phylogenetic relation between the species. The expected correlation matrix for the continuous trait was generated using a Brownian motion^78,79^ (`ape` (version 5.4), using function `corBrownian`) on the mammalian phylogeny from Bininda-Emonds et al. (2007)^74^ adding the missing species (Fig 1a). The full model was compared to a null model using the likelihood ratio test (LRT). Residual R^2^ values were calculated using `R2.resid` function from R package RR2 (version 1.0.2).

### Massively Parallel Reporter Assay (MPRA)

#### MPRA library design

A total of 351 potential *TRNP1* CRE sequences were identified as outlined before. Based on these, the Massively Parallel Reporter Assay (MPRA) oligos were designed as 94mers, where larger sequences were covered by sliding window by 40 bases, resulting in 4,950 oligonucleotide sequences, that are flanked by upstream and downstream priming sites and KpnI/Xbal restriction cut sites as in the original publication^80^. Barcode tag sequences were designed so that they contain all four nucleotides at least once, do not contain stretches of four identical nucleotides, do not contain microRNA seed sequences (retrieved from microRNA Bioconductor R package version 1.28.0) and do not contain restriction cut site sequences for KpnI nor Xbal. The full library of designed oligonucleotides can be found on github (see Data and Code Availability)

#### MPRA library construction

We modified the original MPRA protocol^80^ by using a lentiviral delivery system as previously described^81^, introducing green fluorescent protein (GFP) instead of nano luciferase and changing the sequencing library preparation strategy. In brief oligonucleotide sequences (Custom Array) were amplified using emulsion PCR (Micellula Kit, roboklon) and introduced into the pMPRA plasmid as described previously. The nanoluciferase sequence used in the original publication was replaced by EGFP using Gibson cloning and subsequent insertion into the enhancer library using restriction enzyme digest as in the original publication. Using SFiI the assembled library was transferred into a suitable lentiviral vector (pMPRAlenti1,Addgene #61600). Primer sequences and plasmids used in the MPRA can be found in the analysis github (See Data and Code Availability). To ensure maximum library complexity, transformations that involved the CRE library were performed using electroporation (NEB 10-beta electrocompetent *E.coli*), in all other cloning steps chemically competent *E. coli* (NEB 5-alpha) were used.

Lentiviral particles were produced according to standard methods in HEK 293T cells^82^. The MPRA library was co-transfected with third generation lentiviral plasmids (pMDL-g/pRRE, pRSV-Rev, pMD2.G; Addgene #12251, #12253 #12259) using Lipofectamine 3000. The lentiviral particle containing supernatant was harvested 48 hrs post transfection and filtered using 0.45 μm PES syringe filters. Viral titer was determined by infecting Neuro-2A cells (ATCC CCL-131) and counting GFP positive cells. To this end, N2A cells were infected with a 50/50 volume ratio of viral supernatant to cell suspension with addition of 8 μg/ml Polybrene. Cells were exposed to the lentiviral particles for 24 hrs until medium was exchanged. Selection was performed using Blasticidin starting 48 hrs after infection.

#### MPRA lentiviral transduction

The transduction of the MPRA library was performed in triplicates on two *Homo sapiens* and one *Macaca fascicularis* NPC lines generated as described previously^39^. 2.5 × 10^5^NPCs per line and replicate were dissociated, dissolved in 500 μl cell culture medium containing 8 μg/ml Polybrene and incubated with virus at MOI 12.7 for 1 h at 37°C in suspension^83^. Thereafter cells were seeded on Geltrex and cultured as described above. Virus containing medium was replaced the next day and cells were cultured for additional 24 hrs. Cells were collected, lysed in 100 μl TRI reagent and frozen at −80°C.

#### MPRA sequencing library generation

As input control for RNA expression, DNA amplicon libraries were constructed using 100 - 500 pg plasmid DNA. Library preparation was performed in two successive PCRs. A first PCR introduced the 5’ transposase mosaic end using overhang primers, this was used in the second PCR (Index PCR) to add a library specific index sequence and Illumina Flow Cell adapters. The Adapter PCR was performed in triplicates using DreamTaq polymerase (Thermo Fisher Scientific). Subsequently 1-5 ng of the Adapter PCR product were subjected to the Index PCR using Q5 polymerase. Total RNA from NPCs was extracted using the Direct-zol RNA Microprep Kit (Zymo Research). 500ng of RNA were subjected to reverse transcription using Maxima H Minus RT (Thermo Fisher Scientific) with oligo-dT primers. 50 ng of cDNA were used for library preparation and processed as described for plasmid DNA. Plasmid and cDNA libraries were pooled and quality was evaluated using capillary gel electrophoresis (Agilent Bioanalyzer 2100). Sequencing was performed on an Illumina HiSeq 1500 instrument using a single-index, 50bp, paired-end protocol.

#### MPRA data processing and analysis

MPRA reads were demultiplexed with deML^69^ using i5 and i7 adapter indices from Illumina. Next, we removed barcodes with low sequence quality, requiring a minimum Phred quality score of 10 for all bases of the barcode (zUMIs, `fqfilter.pl` script^84^). Furthermore, we removed reads that had mismatches to the constant region (the first 20 bases of the GFP sequence TCTAGAGTCGCGGCCTTACT). The remaining reads that matched one of the known CRE-tile barcodes were tallied up resulting in a count table. Next, we filtered out CRE tiles that had been detected in only one of the 3 input plasmid library replicates (4202/4950). Counts per million (CPM) were calculated per CRE tile per library (median counts: ~ 900k range: 590k-1,050k). Macaque replicate 3 was excluded due its unusually low correlation with the other samples (Pearson’s r: 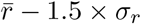). The final regulatory activity for each CRE tile per cell line was calculated as:

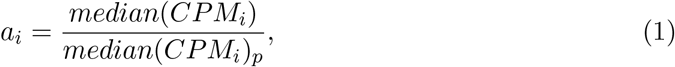

where *a* is regulatory activity, *i* indicates CRE tile and p is the input plasmid library. Median was calculated across the replicates from each cell line.

Given that each tile was overlapping with two other tiles upstream and two downstream, we calculated the total regulatory activity per CRE region in a coverage-sensitive manner, i.e. for each position in the original sequence, mean per-bp-activity across the detected tiles covering it was calculated. The final CRE region activity is the sum across all base positions.

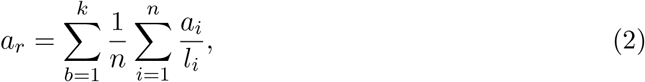

where a_r_ is regulatory activity of CRE region *r*, *b* = 1,…, *k* is the base position of region *r*, *i*,…,*n* are tiles overlapping the position *b*, *a_i_* is tile activity from equation 1 and *l_i_* is tile length. CRE activity and brain phenotypes were associated with one another using PGLS analysis (see above). The number of species varied for each phenotype-CRE pair (brain size: min. 37 for exon1, max. 48 for intron and downstream regions; GI: min. 32 for exon2, max. 37 for intron), therefore the activity of each of the seven CRE regions was used separately to predict either GI or brain size of the respective species.

#### Combining protein evolution rates and intron activity to predict GI across catharrines

PGLS model fits were compared either including only *ω* of TRNP1 protein from Coevol^36^ or including *ω* and intron CRE activity as predictors. For this, the standardised values of either measurement were used, calculated as 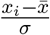, where *x_i_* is each observed value, 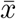 is the mean and *σ* is the standard deviation.

### Transcription Factor analysis

#### RNA-seq library generation

RNA sequencing was performed using the prime-seq method, a bulk derivative of the single cell RNA-seq method SCRB-seq^85^. The full prime-seq protocol including primer sequences can be found at protocols.io (https://www.protocols.io/view/prime-seq-s9veh66). Here we used 10 ng of the isolated RNA from the MPRA experiment and subjected it to the prime-seq protocol. Sequencing was performed on an Illumina HiSeq 1500 instrument with the following setup: read 1 16 bases, read 2 50 bases and i7 index read 8 bases.

#### RNA-seq data processing

Bulk RNA-seq data was generated from the same 9 samples (3 cell lines, 3 biological replicates each) that were assayed in the MPRA. Raw read fastq files were pre-processed using zUMIs (version 2.4.5b)^84^ together with STAR (version STAR_2.6.1c)^86^ to generate expression count tables for barcoded UMI data. Reads were mapped to human reference genome (hg38, Ensembl annotation GRCh38.84). Further filtering was applied keeping genes that were detected in at least 7/9 samples and had on average more than 7 counts, resulting in 17,306 genes. For further analysis, we used normalised and variance stabilised expression estimates as provided by DESeq2^87^.

#### TFBS motif analysis on the intron CRE sequence

TF Position Frequency Matrices (PFM) were retrieved from JASPAR CORE 2020^88^, including only non-redundant vertebrate motifs (746 in total). These were filtered for the expression in our NPC RNA-seq data, leaving 392 TFs with 462 motifs in total.

A Hidden Markov Model (HMM)-based program `Cluster-Buster`^44^ (compiled on Jun 13 2019) was used to infer the enriched TF binding motifs on the intron sequence. First, the auxiliary program `Cluster-Trainer` was used to find the optimal gap parameter between motifs of the same cluster and to obtain weights for each TF based on their motif abundance per kb across catharrine intron CREs from 10 species with available GI measurements. Weights for each motif suggested by `Cluster-Trainer` were supplied to `Cluster-Buster` that we used to find clusters of regulatory binding sites and to infer the enrichment score for each motif on each intron sequence. The program was run with the following parameters: `-g3 -c5 -m3`. To identify the most likely regulators of *TRNP1* that bind to its intron sequence and might influence the evolution of gyrification, we filtered for the motifs that were most abundant across the intron sequences (`Cluster-Trainer` weights >1). These motifs were distinct from one another (mean pairwise distance 0.72). Gene-set enrichment analysis contrasting the TFs with the highest binding potential with the other expressed TFs was conducted using the Bioconductor-package topGO^89^(version 2.40.0) (Table 12).

PGLS model was applied as previously described, using `Cluster-Buster` binding scores across catharrine intron CRE sequences as predictors and predicting either intron activity or GI from the respective species. The relevance of the three TFs that were associated with intron activity was then tested using an additive model and comparing the model likelihoods with reduced models where either of these were dropped.

### Quantification and Statistical Analysis

Data visualizations and statistical analysis was performed using R (version 4.0)^90^. Details of the statistical tests performed in this study can be found in the main text as well as the method details section and supplementary tables. For display Items all relevant parameters like sample size (n), type of statistical test, significance thresholds, degrees of freedom as well as standard deviations can be found in the figure legends.

## Resource Availability

### Lead contact

Further information and requests for resources and reagents should be directed to and will be fulfilled by the lead contact, Wolfgang Enard.

### Materials Availability

Plasmids and cell lines used in this work will be available upon request.

### Data and Code Availability

The RNA-seq data used in this manuscript have been submitted to Array Express (https://www.ebi.ac.uk/arrayexpress/) under the accession number E-MTAB-9951. The MPRA data have been submitted to Array Express under accession number E-MTAB-9952. Additional primate sequences for TRNP1 have been submitted to GenBank (https://www.ncbi.nlm.nih.gov/genbank/) under the accession numbers MW373535 - MW373709.

A compendium containing processing scripts and detailed instructions to reproduce the analysis for this manuscript is available from the following GitHub repository: https://github.com/Hellmann-Lab/Co-evolution-TRNP1-and-GI.

## Author Contributions

M.G. proposed the project and W.E and I.H. conceived the approaches of this study. B.V. designed all initial sequence acquisitions. L.W., M.H., D.R. and J.R. conducted the MPRA assay. M.E. designed and conducted the proliferation assay. J.R., J.G. and M.O. were responsible for all primate cell culture work. Z.K. collected, integrated and analysed all data. M.G. and M.E. provided expertise on Trnp1 function throughout the study. W.E. and I.H. supervised the work and provided guidance in data analysis. Z.K., I.H., and W.E. wrote the manuscript. All authors read, corrected and approved the final manuscript.

## Acknowledgements

This work was supported by the Deutsche Forschungsgemeinschaft (DFG) through LMUex-cellent, SFB1243 (Subproject A14/A15 to W.E. and I.H., respectively), DFG grant HE 7669/1-1 (to I.H.) and the advanced ERC grants ChroNeuroRepair and NeuroCentro (to M.G.) and the Cyliax foundation (to W.E.). We want to thank Christian Roos from the German Primate Center for providing genomic DNA from primates, Deeksha for providing Ferret fibroblast cells, project students Gunnar Kuut and Fatih Sarigoel for helping to generate TRNP1 orthologous sequences, Nikola Vukovi*ć* for helping to establish the MPRA assay, Nika Foglar and Reza Rifat for helping with the proliferation assays and Christoph Neumayr, Tamina Dietl for helping in data analysis.

## Key Resources

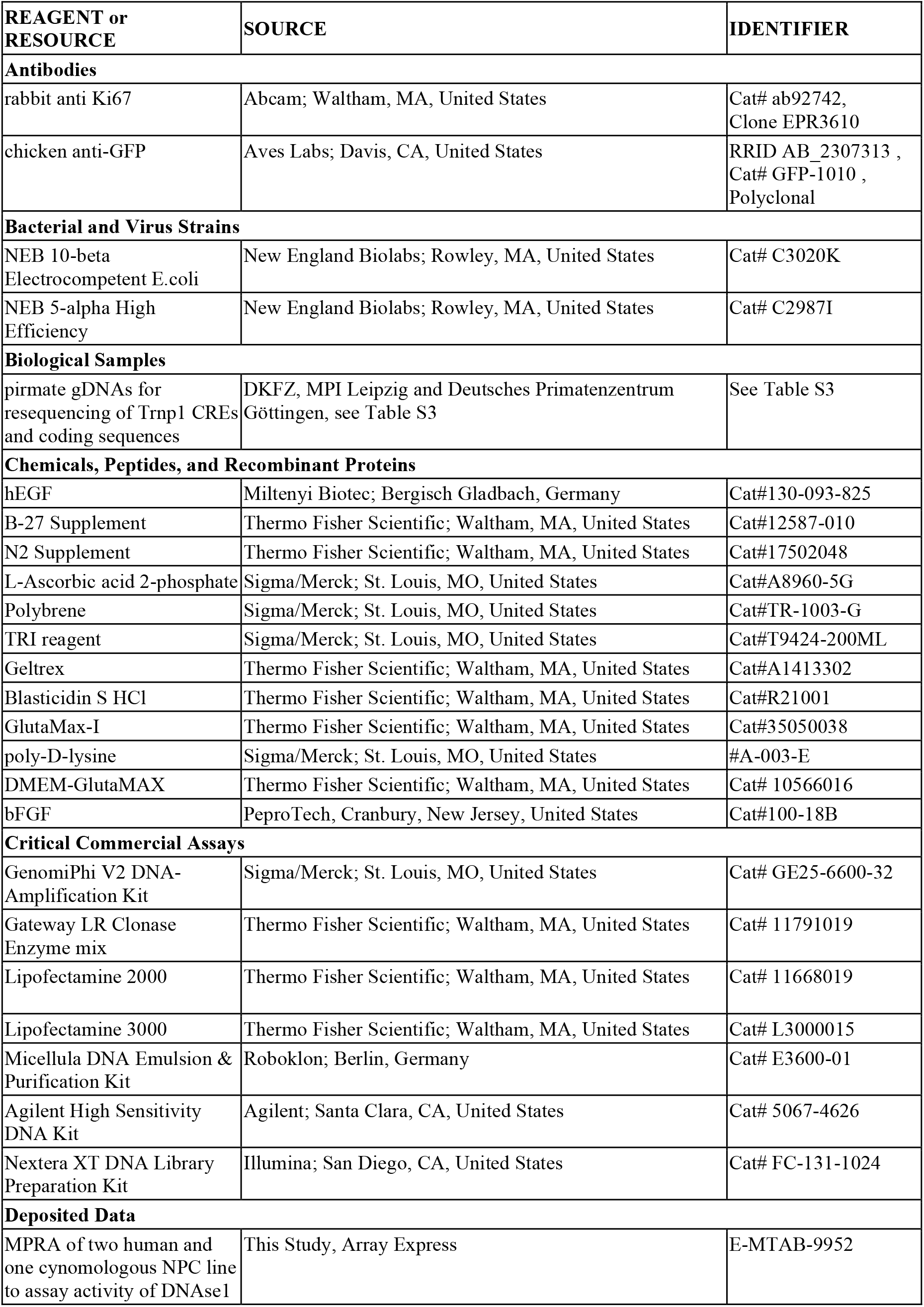
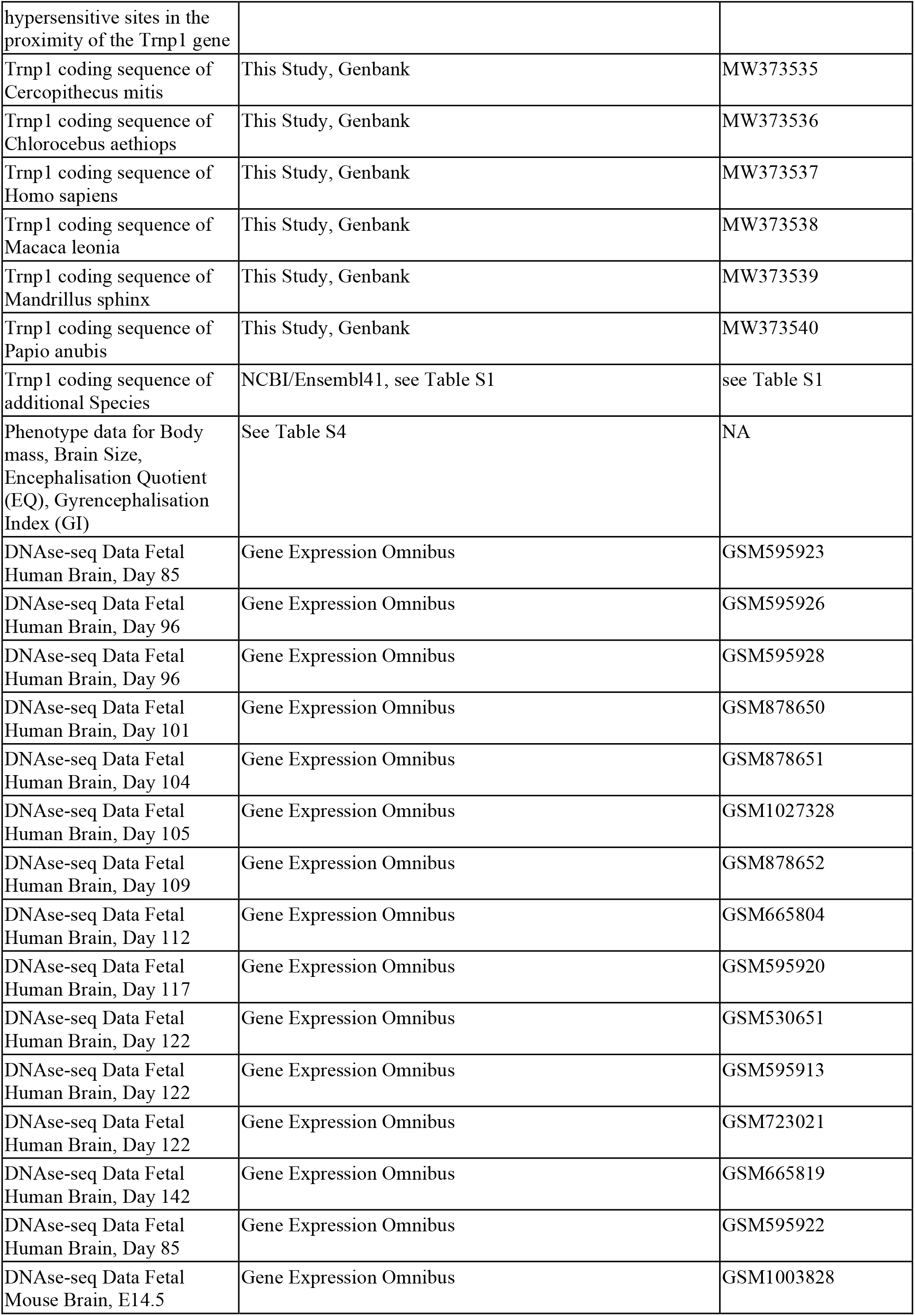
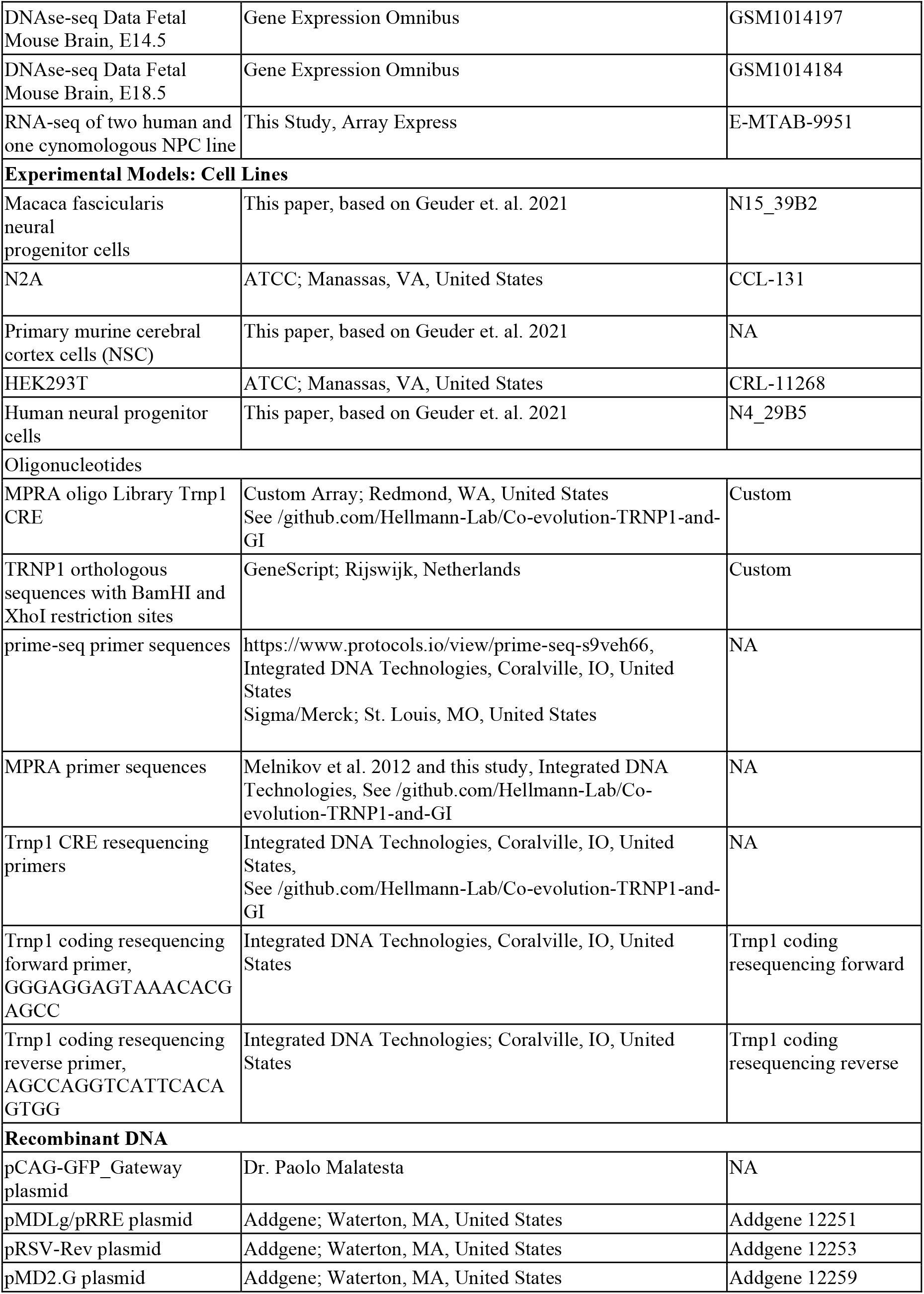
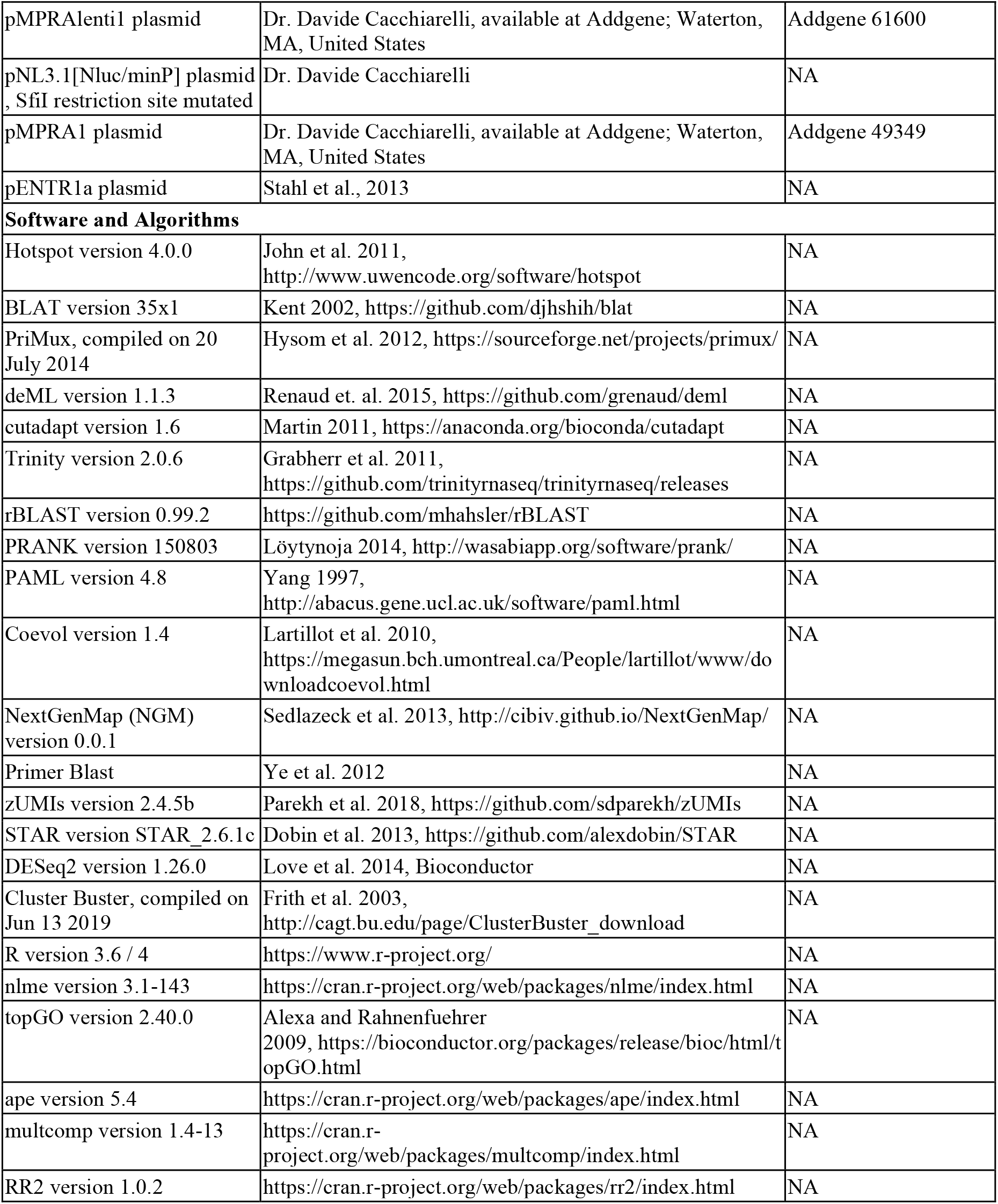

## Supplementary Information

**Supplementary Figure 1.**
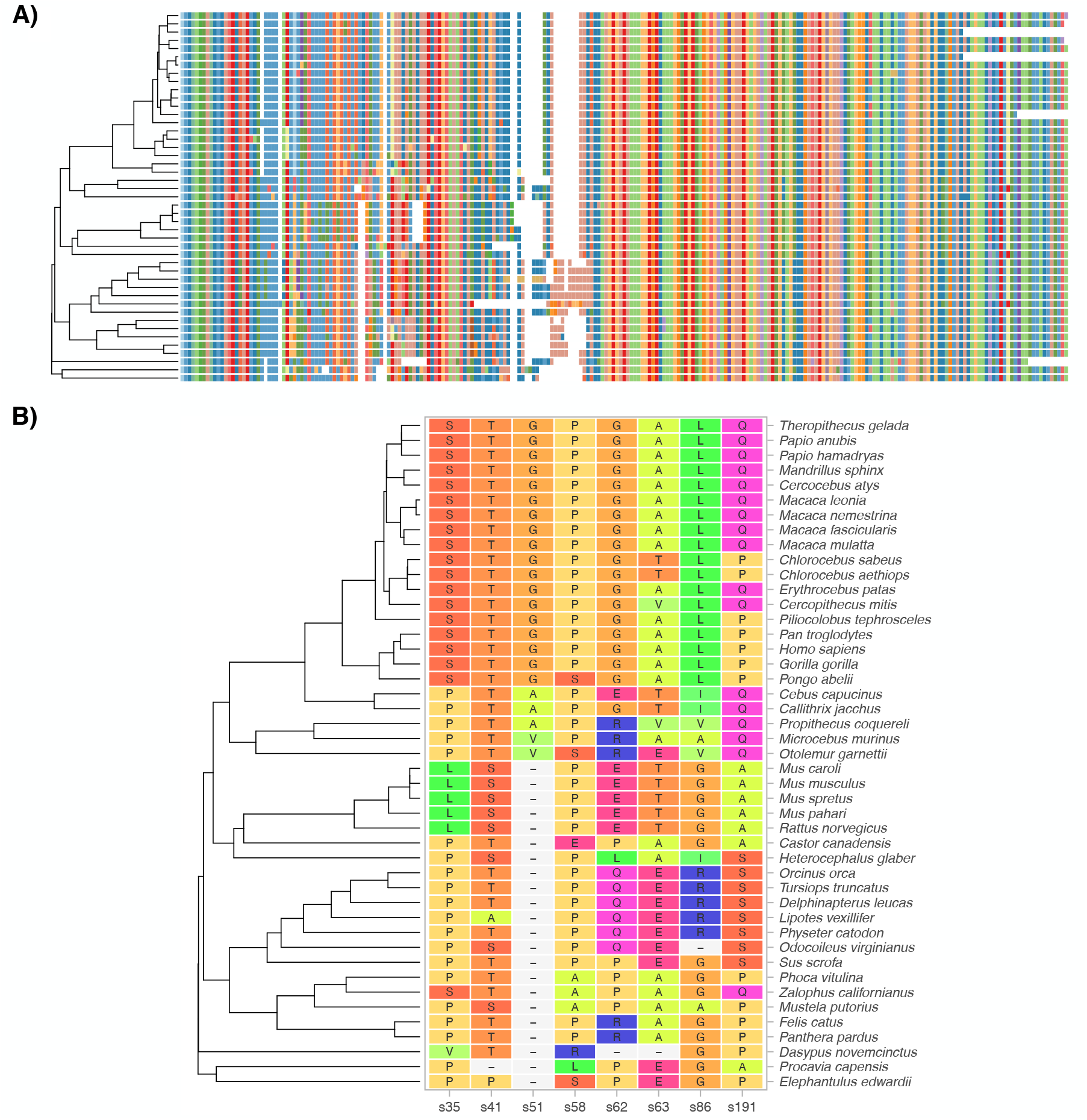
(related to Figure 1)TRNP1 protein-coding sequence analysis. A) Multiple alignment of 45 TRNP1 coding sequences (99.0% completeness) using phylogeny-aware aligner PRANK ([29]). The alignment is 735 bases long, which translates to 245 amino acids (AAs). For comparison: human TRNP1 coding sequence is 227 AA long, whereas mouse - 223 AAs. B) Sites under positive selection across the phylogenetic tree according to PAML ([30]) M8 model (in total 9.8% of sites with *ω* > 1, LRT, *p*-value< 0.001). The depicted sites had a posterior probability Pr(*ω* > 1) > 0.95 according to Naive Empirical Bayes analysis. Colours of the amino acids indicate their relatedness in biochemical properties. Sites with light-grey background and a dash indicate indels.

**Supplementary Figure 2.**
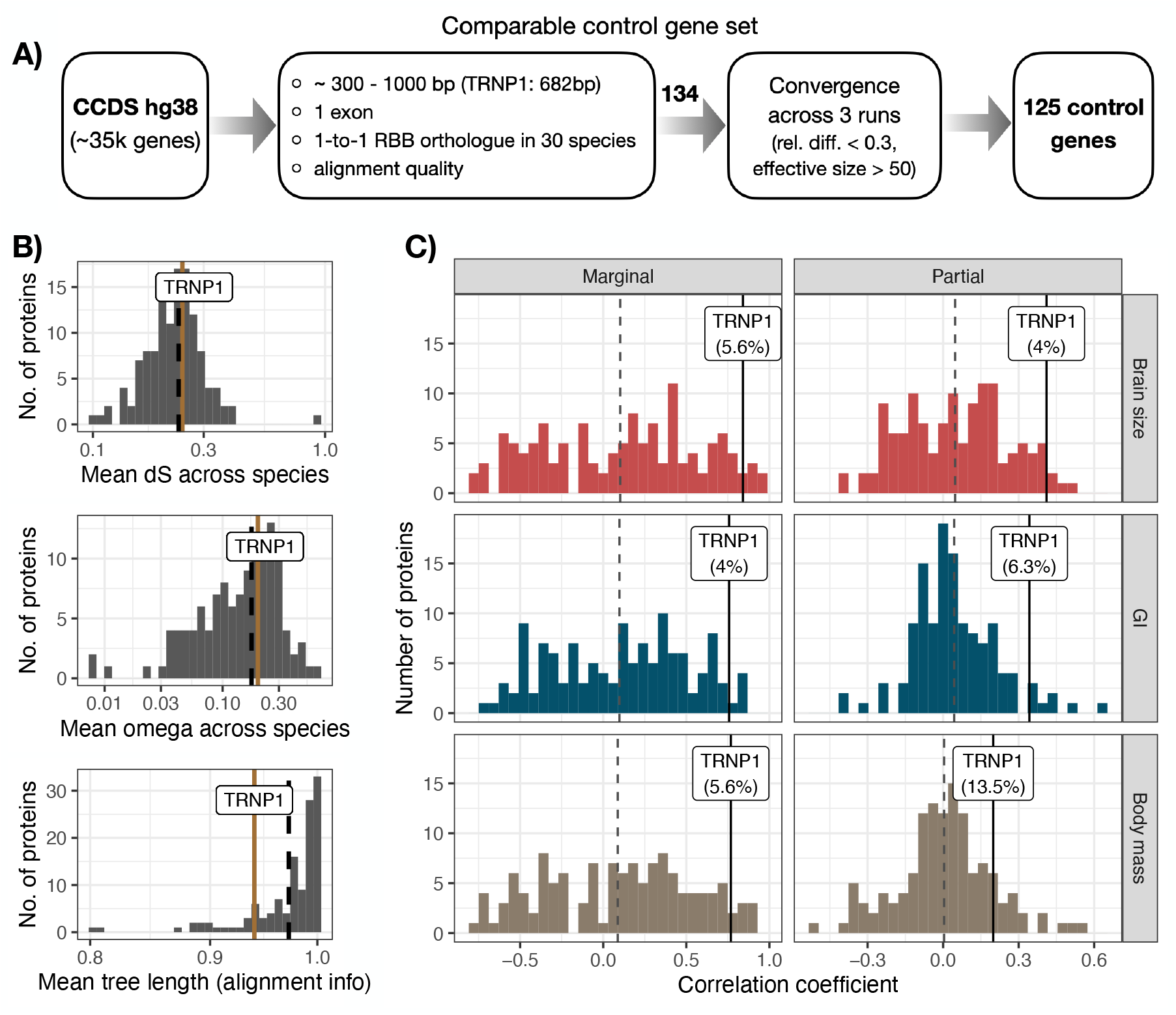
(related to Figure 1) Control protein evolution rate correlation with brain size, gyrification and body mass. A) A flowchart depicting the selection of comparable control proteins to infer the average protein correlation rate with the included phenotypes across the mammalian phylogeny of 30 species. RBB - reciprocal best blat. B) TRNP1 and the control proteins show comparable mutation rates (dS) and protein evolution rates (omega). In addition, all included proteins have high-quality full length alignments (bottom), quantified as the mean relative tree length across all alignment sites per protein. Brown lines indicate TRNP1, black dashed lines - the average across all proteins. C) Marginal and partial correlation distribution of 126 proteins, including TRNP1, with the three phenotypes: brain size, gyrification (GI) and body mass inferred using Coevol ([36]) (combined model). Dashed lines indicate the average across all proteins.

**Supplementary Figure 3.**
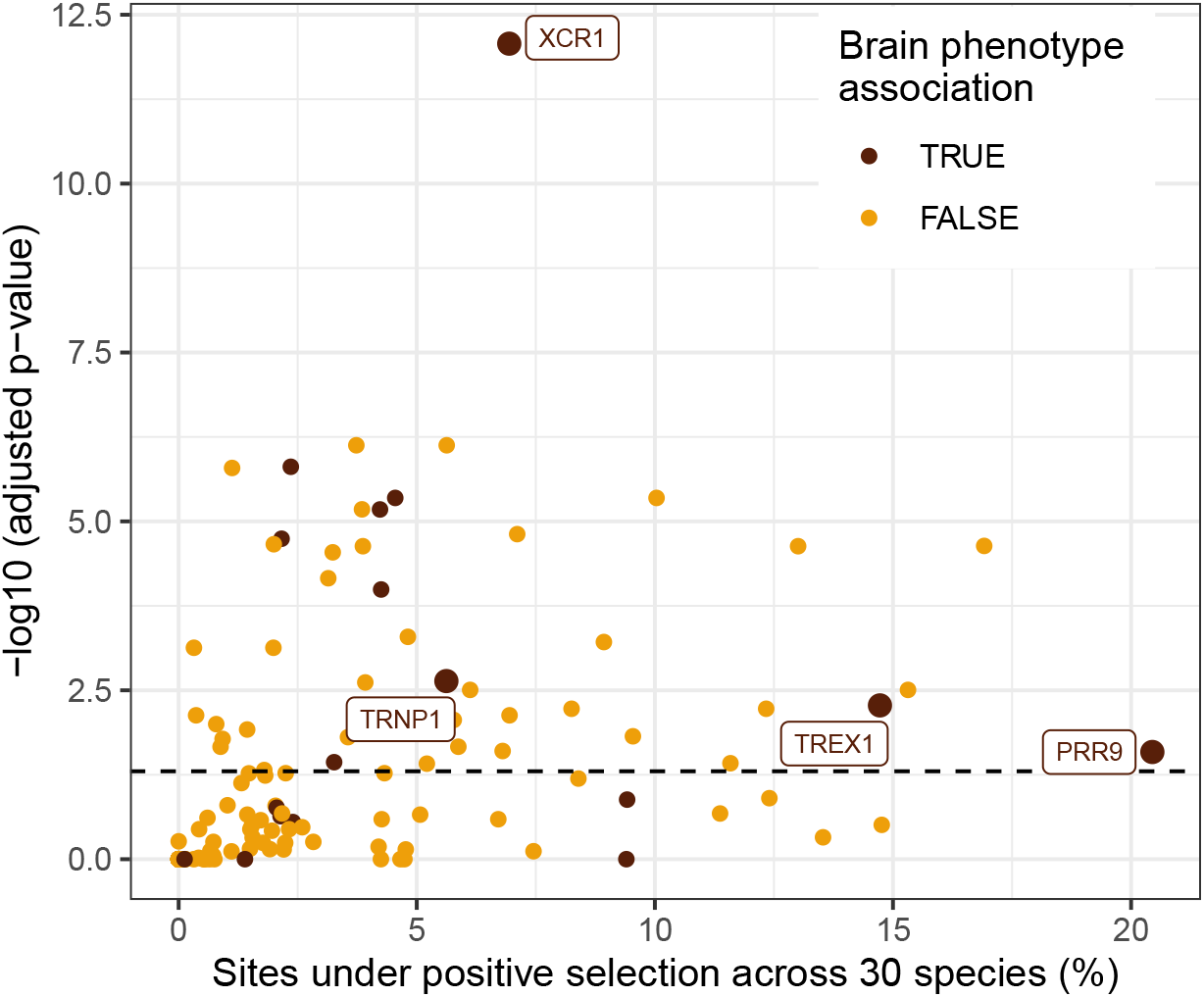
(related to Figure 1) Percentage of sites under positive selection in TRNP1 and control proteins across the 30 species with phenotype information inferred using PAML ([30]). Proteins above the dashed line show a significant proportion of sites under positive selection (*ω* > 1, M8 vs. M7 model, Benjamini-Hochberg adjusted *χ*^2^ *p*-value< 0.05, df=2). Proteins depicted in dark-red color are significantly associated with either brain size or gyrification (Coevol ([36]) combined-model, marginal *pp* <0.95). Only 3 other brain-associated proteins showed significant and higher proportion of sites evolving under positive selection across the tree.

**Supplementary Figure 4.**
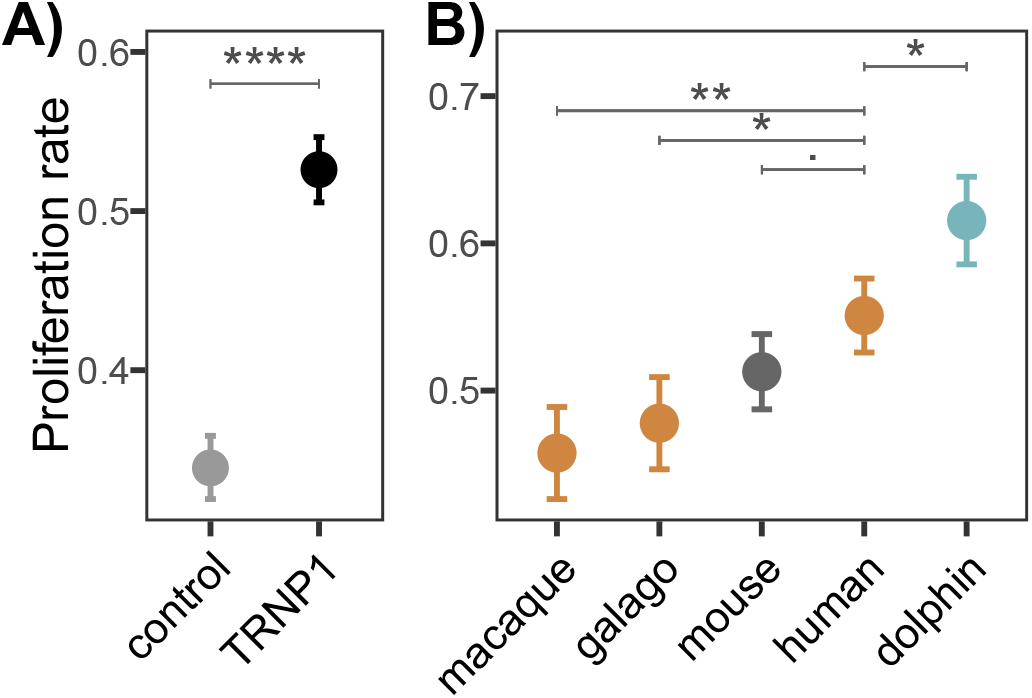
(related to Figure 2) Induced proliferation by TRNP1. A) Proliferation induced in NSCs transfected with TRNP1 (all TRNP1 orthologues combined) compared to the control NSCs transfected only with GFP. TRNP1 presence in NSCs significantly increases the proliferation rates (TRNP1: 0.53 (±0.02), control: 0.34 (±0.02), df=57). B) Proliferation induced in NSCs transfected with TRNP1 orthologous from five different species (Table 8). A), B) Bars indicate standard errors of logistic regression and asterisks indicate the significance of pairwise comparisons (Tukey test, *p*-value: .<0.1, *<0.05, **<0.001, ****<2e-16).

**Supplementary Figure 5.**
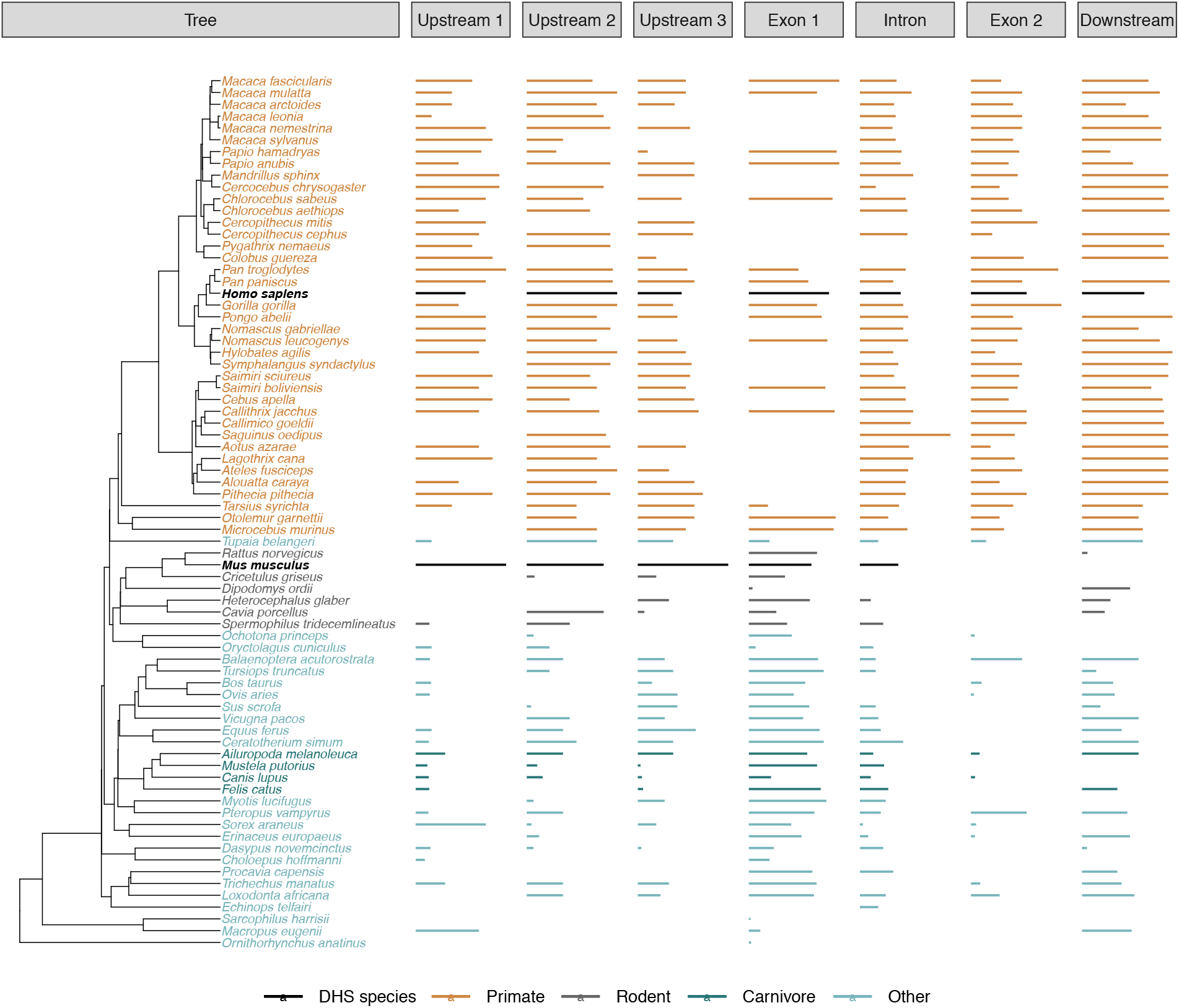
(related to Figure 3) Length of the covered CRE sequences in the MPRA library across the tree. Species for which the regions were inferred based on DNase-Hypersensitive Sites (DHS) from embryonic brain ([38]) are marked in bold and black: human (*Homo sapiens*) and mouse (*Mus musculus*). These species do not show extreme differences in length compared to others (human: 5/7, mouse: 3/5 regions within the 10% and 90% quantiles). The orthologous CRE sequence length differs strongly between primate and non-primate species, being on average 1.8 to 2.8 times longer in the primate species than in the other mammals.

**Supplementary Figure 6.**
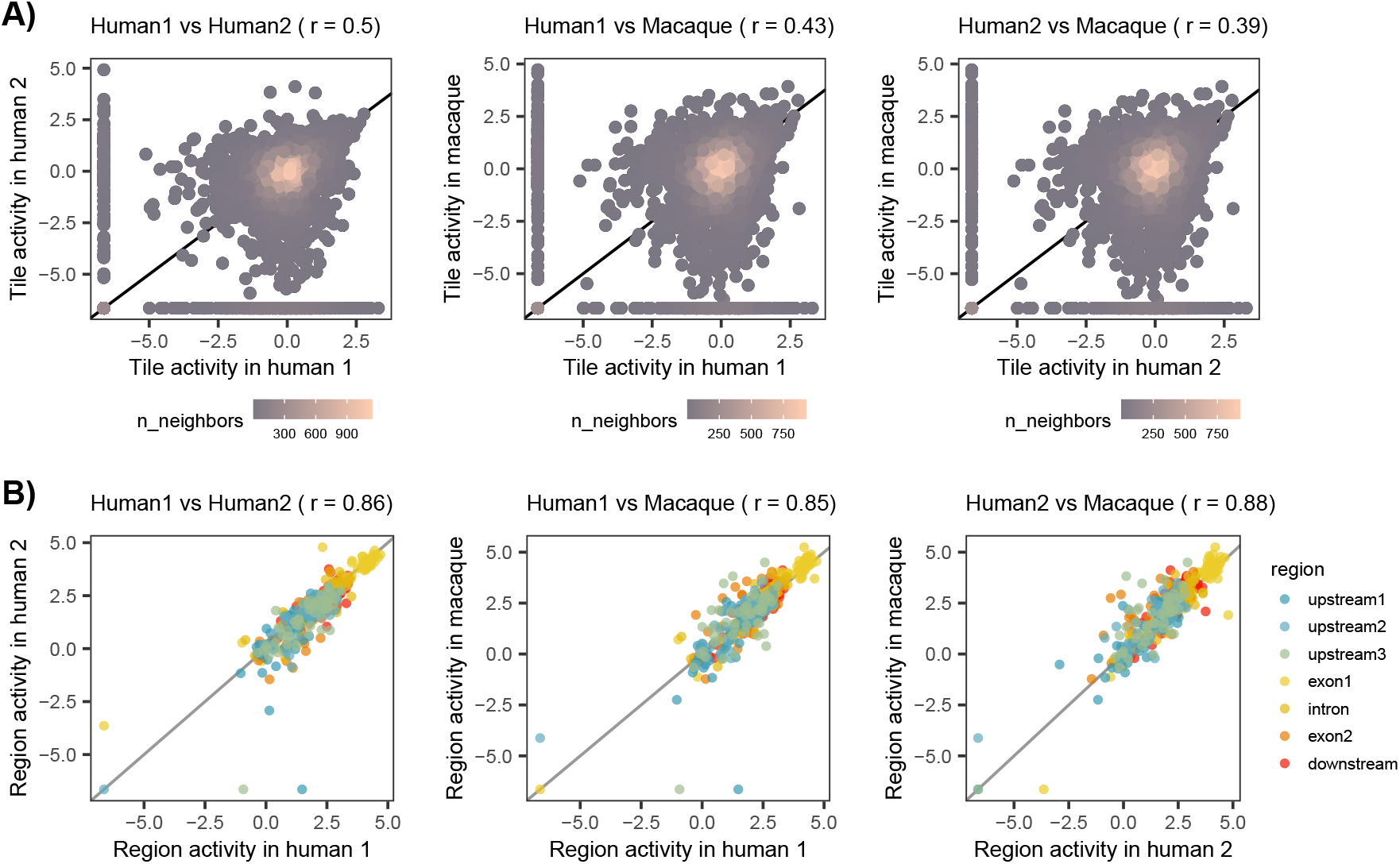
(related to Figure 3) Analysis of massively parallel reporter assay (MPRA) data. A) Pairwise correlation of the log2-transformed CRE tile activity between the three transduced cell lines: human 1, human 2 and macaque. Pearson’s *r* is specified in the brackets of figure titles. B) Pairwise correlation of the log2-transformed summarized activity per CRE region between cell lines. Pearson’s *r* is specified in the brackets of figure titles.

**Supplementary Table 1.**
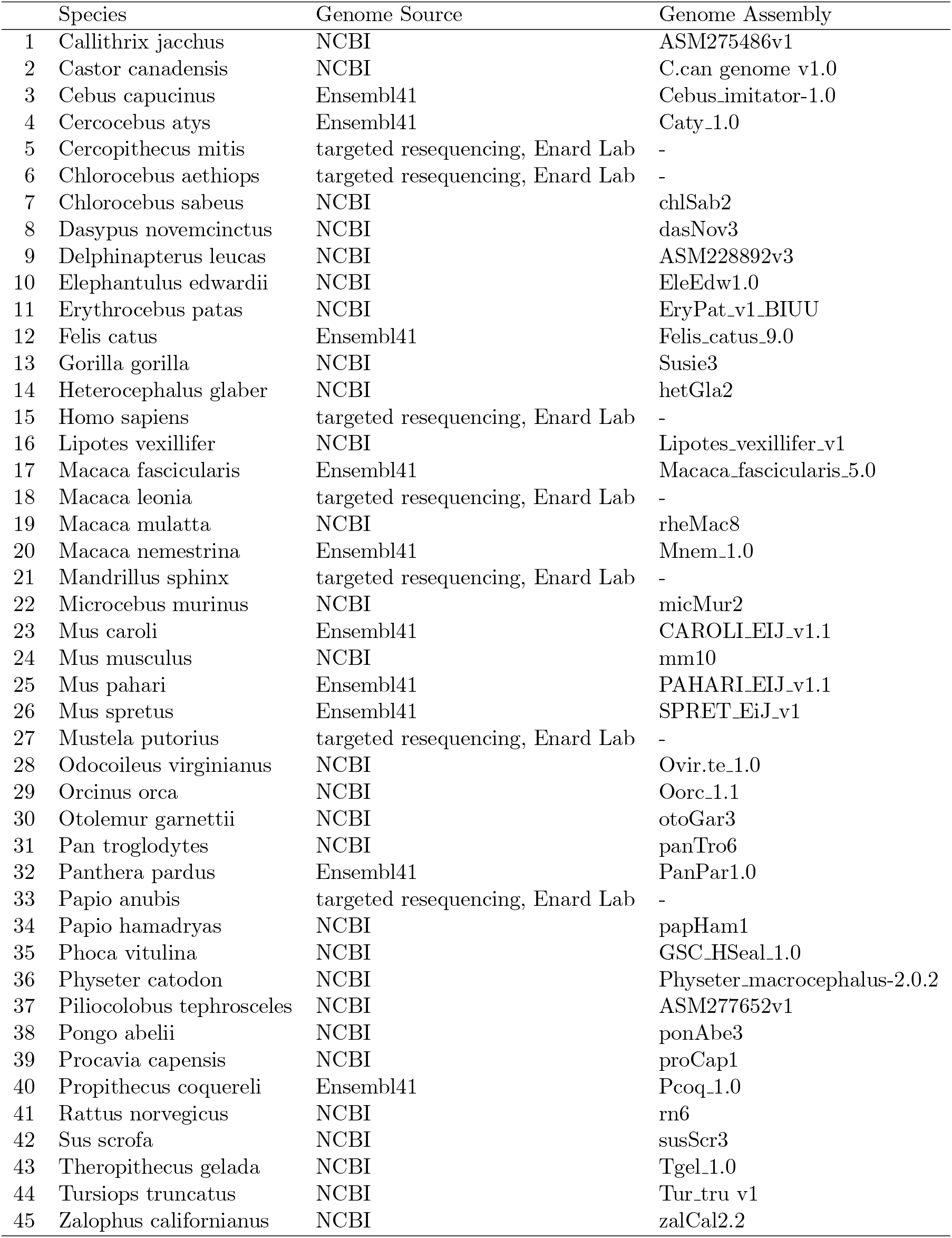
Genome sources of the TRNP1 protein coding sequences

**Supplementary Table 2.** *TRNP1* DNase hypersensitive sites in Human and Mouse

|  | Chromosome | Start | End | ID | Species |
| --- | --- | --- | --- | --- | --- |
| 1 | chr1 | 27293479 | 27293766 | upstream1 | Human |
| 2 | chr1 | 27310581 | 27310877 | upstream2 | Human |
| 3 | chr1 | 27318087 | 27318439 | upstream3 | Human |
| 4 | chr1 | 27319449 | 27321900 | promexon1 | Human |
| 5 | chr1 | 27323922 | 27324667 | intron | Human |
| 6 | chr1 | 27327174 | 27327461 | exon2 | Human |
| 7 | chr1 | 27328171 | 27328804 | downstream | Human |
| 8 | chr4 | 133494338 | 133494835 | intron | Mouse |
| 9 | chr4 | 133495742 | 133496109 | unique1 | Mouse |
| 10 | chr4 | 133497135 | 133498824 | promexon1 | Mouse |
| 11 | chr4 | 133504292 | 133504667 | unique2 | Mouse |
| 12 | chr4 | 133504895 | 133505292 | unique3 | Mouse |
| 13 | chr4 | 133508796 | 133509525 | upstream3 | Mouse |
| 14 | chr4 | 133511090 | 133511417 | upstream2 | Mouse |
| 15 | chr4 | 133512990 | 133513416 | upstream1 | Mouse |

**Supplementary Table 3.**
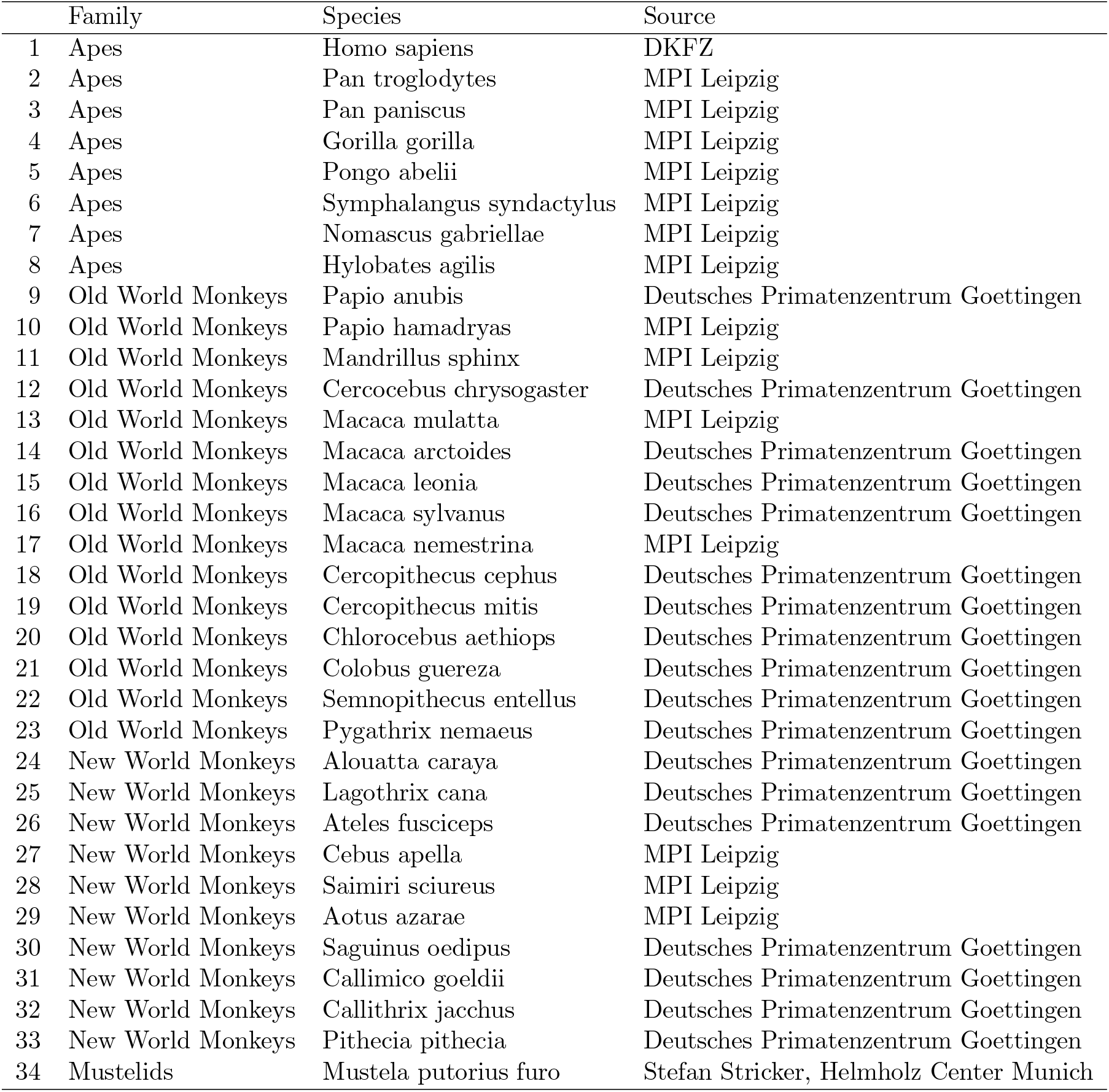
gDNA samples

**Supplementary Table 4.**
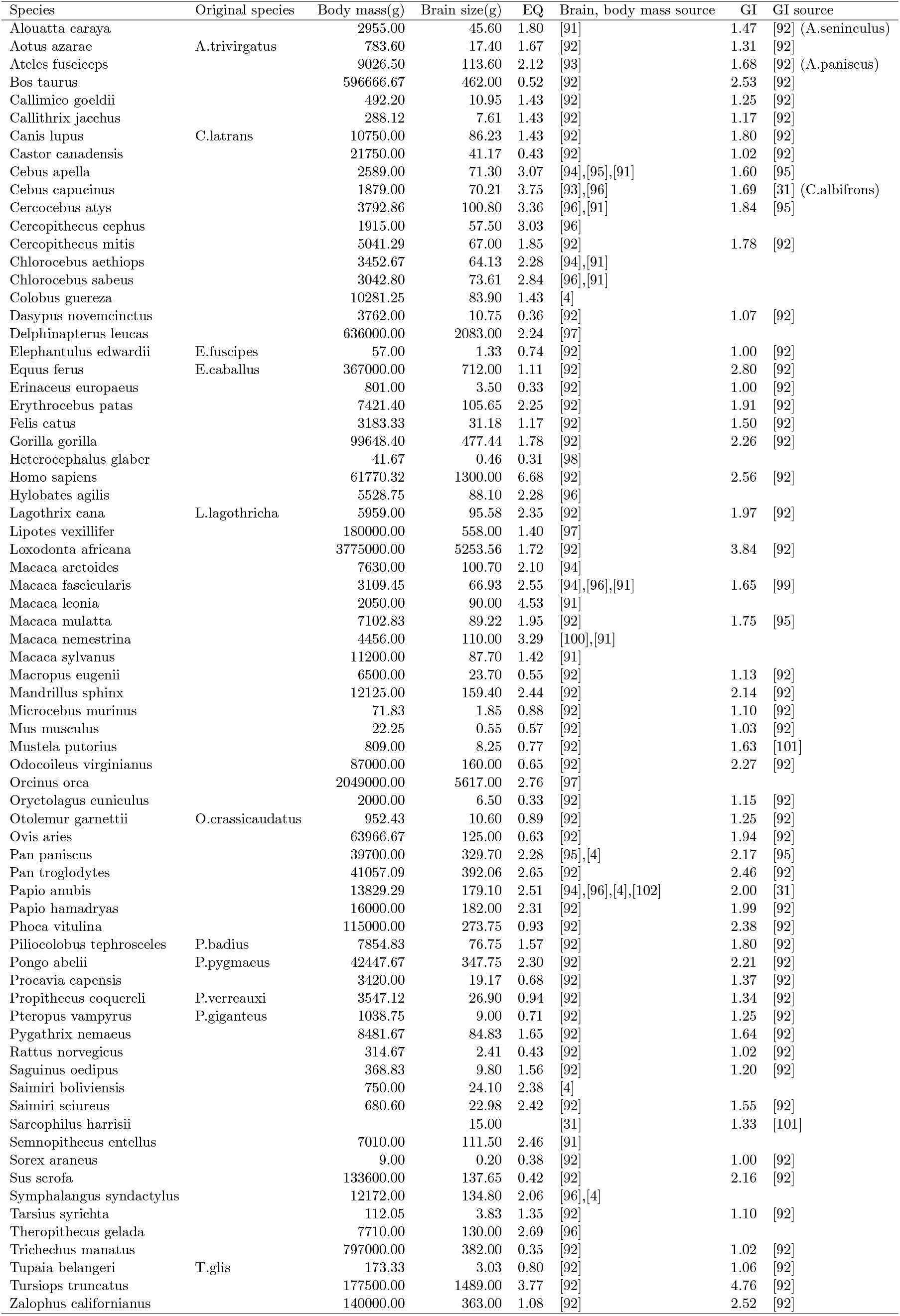
Phenotype data and its source publications used in this study. In cases where there are multiple sources listed, mean across the individual measurements was calculated. For 11 species, the phenotype data of their close sister species (column ‘‘Original species”) was used. For 3 additional species with only missing GI, this information was borrowed from the indicated sister species (column “GI source” in the brackets)

**Supplementary Table 5.** TRNP1 amino acid sites under positive selection across the phylogeny according to Naive Empirical Bayes analysis [30] (*: P>95%; **: P>99%)

| Alignment position | Pr($\omega > 1$) | post mean $\omega$ |
| --- | --- | --- |
| 35 | 0.985* | 1.04 |
| 41 | 0.995** | 1.05 |
| 51 | 0.952* | 1.01 |
| 58 | 0.967* | 1.02 |
| 62 | 0.995** | 1.05 |
| 63 | 0.984* | 1.04 |
| 86 | 0.991** | 1.04 |
| 191 | 0.980* | 1.04 |

**Supplementary Table 6.** Pairwise correlations between the substitution rates of TRNP1 (dS: synonymous substitution rates, *ω*: the ratio of the non-synonymous over the synonymous substitution rates) and the rate of change in either GI, brain size or body mass estimated either separately across 30 mammalian species or together using Coevol ([36]). Partial correlations are the maximally controlled correlations, controlling for all other included variables

| Parameter 1 | Parameter 2 | Marginal Correlation<br>(Posterior Probability) | Partial Correlation<br>(Posterior Probability) | Type of Estimate |
| --- | --- | --- | --- | --- |
| brain mass | $\omega$ | 0.789 (0.97) | 0.744 (0.94) | Separate |
| brain mass | dS | -0.554 (0.99) | 0.012 (0.5) | Separate |
| $\omega$ | dS | -0.599 (0.93) | -0.309 (0.69) | Separate |
| GI | $\omega$ | 0.769 (0.98) | 0.775 (0.96) | Separate |
| GI | dS | -0.42 (0.96) | 0.093 (0.57) | Separate |
| $\omega$ | dS | -0.525 (0.91) | -0.335 (0.71) | Separate |
| body mass | $\omega$ | 0.696 (0.95) | 0.677 (0.91) | Separate |
| body mass | dS | -0.416 (0.94) | 0.068 (0.54) | Separate |
| $\omega$ | dS | -0.53 (0.89) | -0.364 (0.73) | Separate |
| brain mass | $\omega$ | 0.827 (0.97) | 0.403 (0.83) | Combined |
| body mass | $\omega$ | 0.755 (0.97) | 0.19 (0.65) | Combined |
| GI | $\omega$ | 0.748 (0.98) | 0.341 (0.77) | Combined |
| body mass | dS | -0.459 (0.96) | 0.266 (0.78) | Combined |
| GI | dS | -0.466 (0.97) | 0.17 (0.69) | Combined |
| $\omega$ | dS | -0.537 (0.94) | -0.134 (0.61) | Combined |
| brain mass | dS | -0.61 (0.99) | -0.321 (0.8) | Combined |
| GI | brain mass | 0.811 (1) | 0.306 (0.8) | Combined |
| brain mass | body mass | 0.905 (1) | 0.531 (0.9) | Combined |
| GI | body mass | 0.679 (1) | -0.239 (0.8) | Combined |

**Supplementary Table 7.** Model selection results between logistic regression models that predict the proportion of proliferating mouse NSCs in the presence of TRNP1 orthologues or GFP control (LRT). n=donor mouse (batch). Proliferation is best predicted by the presence of TRNP1 together with the donor mouse to correct for the batch

| Model | Resid. Df | Resid. Dev | Df | Deviance | Pr(>Chi) | Test |
| --- | --- | --- | --- | --- | --- | --- |
| 0: prolif(%) ∼ 1 | 56 | 1312.80 | | | | TRNP1 vs. ctrl |
| 1: prolif(%) ∼ TRNP1 | 55 | 1058.00 | 1 | 254.80 | 2.3E-57 | TRNP1 vs. ctrl |
| 2: prolif(%) ∼ TRNP1+n | 44 | 209.80 | 11 | 848.20 | 8.4E-175 | TRNP1 vs. ctrl |
| 0: prolif(%) ∼ 1 | 44 | 867.96 | | | | TRNP1 ortologues |
| 1: prolif(%) ∼ orthologue | 40 | 790.69 | 4 | 77.27 | 6.6E-16 | TRNP1 ortologues |
| 2: prolif(%) ∼ orthologue+n | 29 | 108.85 | 11 | 681.85 | 4.2E-139 | TRNP1 ortologues |

**Supplementary Table 8.** The induced NSC proliferation rates by the different TRNP1 orthologues (according to model 2 from Table 7)

| Species | Proliferation rate | Std. Error |
| --- | --- | --- |
| macaque | 0.46 | 0.031 |
| galago | 0.48 | 0.031 |
| mouse | 0.51 | 0.026 |
| human | 0.55 | 0.025 |
| dolphin | 0.62 | 0.03 |

**Supplementary Table 9.**
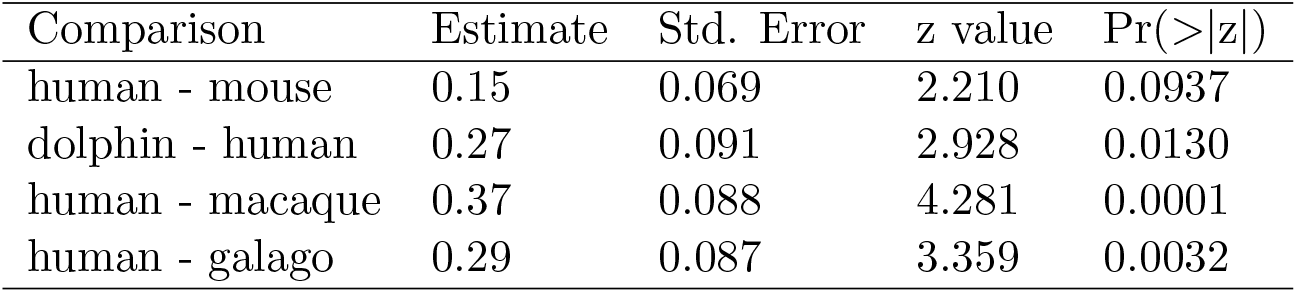
Pairwise proliferation rate comparison between the TRNP1 orthologues of interest (Tukey test)

**Supplementary Table 10.** PGLS model selection using LRT to test whether CRE activity of the 7 *TRNP1* regulatory regions is predictive for either brain size or gyrification (GI) across species. The reduced model contains intercept as the only predictor

| Model | Value | Std.Error | df | L.Ratio | LRT p-value |
| --- | --- | --- | --- | --- | --- |
| log2(brain size) ~ log2(upstream1) | 0.01 | 0.152 | 1 | 0.01 | 0.929 |
| log2(brain size) ~ log2(upstream2) | -0.28 | 0.234 | 1 | 1.43 | 0.232 |
| log2(brain size) ~ log2(upstream3) | -0.30 | 0.223 | 1 | 1.92 | 0.166 |
| log2(brain size) ~ log2(exon1) | 0.31 | 0.487 | 1 | 0.42 | 0.517 |
| log2(brain size) ~ log2(intron) | 0.35 | 0.399 | 1 | 0.78 | 0.377 |
| log2(brain size) ~ log2(exon2) | 0.28 | 0.286 | 1 | 0.97 | 0.324 |
| log2(brain size) ~ log2(downstream) | 0.18 | 0.474 | 1 | 0.16 | 0.693 |
| log2(GI) ~ log2(upstream1) | 0.01 | 0.073 | 1 | 0.01 | 0.929 |
| log2(GI) ~ log2(upstream2) | -0.04 | 0.060 | 1 | 0.53 | 0.468 |
| log2(GI) ~ log2(upstream3) | -0.06 | 0.048 | 1 | 1.51 | 0.219 |
| log2(GI) ~ log2(exon1) | 0.07 | 0.088 | 1 | 0.62 | 0.431 |
| log2(GI) ~ log2(intron) | 0.14 | 0.086 | 1 | 2.75 | 0.097 |
| log2(GI) ~ log2(exon2) | 0.10 | 0.086 | 1 | 1.33 | 0.250 |
| log2(GI) ~ log2(downstream) | 0.03 | 0.114 | 1 | 0.07 | 0.787 |

**Supplementary Table 11.** PGLS model selection using LRT to test whether the association between GI and intron CRE activity on the Old World monkey and great ape branch is consistent across three independent cell lines. The reduced model contains intercept as the only predictor

| Model | Value | Std.Error | df | L.Ratio | LRT p-value | Cell line |
| --- | --- | --- | --- | --- | --- | --- |
| log2(GI) ~ log2(intron) | 0.20 | 0.059 | 1 | 8.66 | 0.003 | human1 |
| log2(GI) ~ log2(intron) | 0.14 | 0.071 | 1 | 3.81 | 0.051 | human2 |
| log2(GI) ~ log2(intron) | 0.10 | 0.059 | 1 | 3.31 | 0.069 | macaque |

**Supplementary Table 12.** Enriched Gene Ontology Terms (Fisher’s *p*-value< 0.05) of the 22 TFs with binding site enrichment on the intron CRE sequences from the 10 catarrhine species. Background: all expressed TFs included in the motif binding enrichment analysis (392)

| GO.ID | Term | Annotated | Significant | Expected | Fisher's P |
| --- | --- | --- | --- | --- | --- |
| GO:0010817 | regulation of hormone levels | 26 | 5 | 1.47 | 0.011 |
| GO:0042127 | regulation of cell population proliferation | 117 | 12 | 6.60 | 0.012 |
| GO:0008285 | negative regulation of cell population proliferation | 61 | 8 | 3.44 | 0.012 |
| GO:0043523 | regulation of neuron apoptotic process | 20 | 4 | 1.13 | 0.020 |
| GO:1901615 | organic hydroxy compound metabolic process | 21 | 4 | 1.18 | 0.024 |
| GO:0051402 | neuron apoptotic process | 23 | 4 | 1.30 | 0.033 |
| GO:1903706 | regulation of hemopoiesis | 47 | 6 | 2.65 | 0.037 |
| GO:0006325 | chromatin organization | 35 | 5 | 1.97 | 0.037 |
| GO:0008283 | cell population proliferation | 135 | 12 | 7.62 | 0.039 |
| GO:0090596 | sensory organ morphogenesis | 36 | 5 | 2.03 | 0.042 |
| GO:0006259 | DNA metabolic process | 25 | 4 | 1.41 | 0.044 |

## References

[1] S. M. Reader, Y. Hager, and K. N. Laland. “The evolution of primate general and cultural intelligence”. en. In: Philosophical transactions of the Royal Society of London. Series B, Biological sciences 366.1567 (2011), pp. 1017–1027. ISSN: 0962-8436, 1471-2970. DOI: 10.1098/rstb.2010.0342.

[2] A. R. DeCasien, R. A. Barton, and J. P. Higham. “Understanding the human brain: insights from comparative biology”. en. In: Trends in cognitive sciences 26.5 (2022), pp. 432–445. ISSN: 1364-6613, 1879-307X. DOI: 10.1016/j.tics.2022.02.003.

[3] S. H. Montgomery, N. I. Mundy, and R. A. Barton. “Brain evolution and development: adaptation, allometry and constraint”. en. In: Proceedings. Biological sciences / The Royal Society 283.1838 (2016). ISSN: 0962-8452, 1471-2954. DOI: 10.1098/rspb.2016.0433.

[4] A. M. Boddy, M. R. McGowen, C. C. Sherwood, L. I. Grossman, M. Goodman, and D. E. Wildman. “Comparative analysis of encephalization in mammals reveals relaxed constraints on anthropoid primate and cetacean brain scaling”. en. In: Journal of evolutionary biology 25.5 (2012), pp. 981–994. ISSN: 1010-061X, 1420-9101. DOI: 10.1111/j.1420-9101.2012.02491.x.

[5] E. Lewitus, I. Kelava, and W. B. Huttner. “Conical expansion of the outer sub-ventricular zone and the role of neocortical folding in evolution and development”. en. In: Frontiers in human neuroscience 7 (2013), p. 424. ISSN: 1662-5161. DOI: 10.3389/fnhum.2013.00424.

[6] J. B. Smaers et al. “The evolution of mammalian brain size”. en. In: Science advances 7.18 (2021). ISSN: 2375-2548. DOI: 10.1126/sciadv.abe2101.

[7] I. Kelava, E. Lewitus, and W. B. Huttner. “The secondary loss of gyrencephaly as an example of evolutionary phenotypical reversal”. en. In: Frontiers in neuroanatomy 7 (2013), p. 16. ISSN: 1662-5129. DOI: 10.3389/fnana.2013.00016.

[8] A. R. DeCasien, S. A. Williams, and J. P. Higham. “Primate brain size is predicted by diet but not sociality”. en. In: Nature ecology & evolution 1.5 (2017), p. 112. ISSN: 2397-334X. DOI: 10.1038/s41559-017-0112.

[9] S. A. Heldstab, K. Isler, S. M. Graber, C. Schuppli, and C. P. van Schaik. “The economics of brain size evolution in vertebrates”. en. In: Current biology: CB 32.12 (2022), R697–R708. ISSN: 0960-9822, 1879-0445. DOI: 10.1016/j.cub.2022.04.096.

[10] A. Pinson and W. B. Huttner. “Neocortex expansion in development and evolution-from genes to progenitor cell biology”. en. In: Current opinion in cell biology 73 (2021), pp. 9–18. ISSN: 0955-0674, 1879-0410. DOI: 10.1016/j.ceb.2021.04.008.

[11] L. Del-Valle-Anton and V. Borrell. “Folding brains: from development to disease modeling”. en. In: Physiological reviews 102.2 (2022), pp. 511–550. ISSN: 0031-9333, 1522-1210. DOI: 10.1152/physrev.00016.2021.

[12] A. Villalba, M. Götz, and V. Borrell. “The regulation of cortical neurogenesis”. en. In: Current topics in developmental biology 142 (2021), pp. 1–66. ISSN: 0070-2153, 1557-8933. DOI: 10.1016/bs.ctdb.2020.10.003.

[13] M. Florio et al. “Human-specific gene ARHGAP11B promotes basal progenitor amplification and neocortex expansion”. en. In: Science 347.6229 (2015), pp. 1465–1470. ISSN: 0036-8075, 1095-9203. DOI: 10.1126/science.aaa1975.

[14] N. Kalebic, C. Gilardi, M. Albert, T. Namba, K. R. Long, M. Kostic, B. Langen, and W. B. Huttner. “Human-specific ARHGAP11B induces hallmarks of neocortical expansion in developing ferret neocortex”. en. In: eLife 7 (2018). ISSN: 2050-084X. DOI: 10.7554/eLife.41241.

[15] M. Heide, C. Haffner, A. Murayama, Y. Kurotaki, H. Shinohara, H. Okano, E. Sasaki, and W. B. Huttner. “Human-specific ARHGAP11B increases size and folding of primate neocortex in the fetal marmoset”. en. In: Science 369.6503 (2020), pp. 546–550. ISSN: 0036-8075, 1095-9203. DOI: 10.1126/science.abb2401.

[16] A. Pinson et al. “Human TKTL1 implies greater neurogenesis in frontal neocortex of modern humans than Neanderthals”. en. In: Science 377.6611 (2022), eabl6422. ISSN: 0036-8075, 1095-9203. DOI: 10.1126/science.abl6422.

[17] I. T. Fiddes et al. “Human-Specific NOTCH2NL Genes Affect Notch Signaling and Cortical Neurogenesis”. en. In: Cell 173.6 (2018), 1356–1369.e22. ISSN: 0092-8674, 1097-4172. DOI: 10.1016/j.cell.2018.03.051.

[18] I. K. Suzuki et al. “Human-Specific NOTCH2NL Genes Expand Cortical Neurogenesis through Delta/Notch Regulation”. en. In: Cell 173.6 (2018), 1370–1384.e16. ISSN: 0092-8674, 1097-4172. DOI: 10.1016/j.cell.2018.03.067.

[19] J. Liu et al. “The Primate-Specific Gene TMEM14B Marks Outer Radial Glia Cells and Promotes Cortical Expansion and Folding”. en. In: Cell stem cell 21.5 (2017), 635–649.e8. ISSN: 1934-5909, 1875-9777. DOI: 10.1016/j.stem.2017.08.013.

[20] X.-C. Ju, Q.-Q. Hou, A.-L. Sheng, K.-Y. Wu, Y. Zhou, Y. Jin, T. Wen, Z. Yang, X. Wang, and Z.-G. Luo. “The hominoid-specific gene TBC1D3 promotes generation of basal neural progenitors and induces cortical folding in mice”. en. In: eLife 5 (2016). ISSN: 2050-084X. DOI: 10.7554/eLife.18197.

[21] J. L. Boyd, S. L. Skove, J. P. Rouanet, L.-J. Pilaz, T. Bepler, R. Gordân, G. A. Wray, and D. L. Silver. “Human-chimpanzee differences in a FZD8 enhancer alter cell-cycle dynamics in the developing neocortex”. en. In: Current biology: CB 25.6 (2015), pp. 772–779. ISSN: 0960-9822, 1879-0445. DOI: 10.1016/j.cub.2015.01.041.

[22] A. M. Boddy, P. W. Harrison, S. H. Montgomery, J. A. Caravas, M. A. Raghanti, K. A. Phillips, N. I. Mundy, and D. E. Wildman. “Evidence of a Conserved Molecular Response to Selection for Increased Brain Size in Primates”. en. In: Genome biology and evolution 9.3 (2017), pp. 700–713. ISSN: 1759-6653. DOI: 10.1093/gbe/evx028.

[23] R. Stahl et al. “Trnp1 regulates expansion and folding of the mammalian cerebral cortex by control of radial glial fate”. en. In: Cell 153.3 (2013), pp. 535–549. ISSN: 0092-8674, 1097-4172. DOI: 10.1016/j.cell.2013.03.027.

[24] G.-A. Pilz et al. “Amplification of progenitors in the mammalian telencephalon includes a new radial glial cell type”. en. In: Nature communications 4 (2013), p. 2125. ISSN: 2041-1723. DOI: 10.1038/ncomms3125.

[25] C. Kerimoglu et al. “H3 acetylation selectively promotes basal progenitor proliferation and neocortex expansion”. en. In: Science advances 7.38 (2021), eabc6792. ISSN: 2375-2548. DOI: 10.1126/sciadv.abc6792.

[26] M. Á. Martínez-Martínez, C. De Juan Romero, V. Fernández, A. Cárdenas, M. Götz, and V. Borrell. “A restricted period for formation of outer subventricular zone defined by Cdh1 and Trnp1 levels”. en. In: Nature communications 7 (2016), p. 11812. ISSN: 2041-1723. DOI: 10.1038/ncomms11812.

[27] M. Esgleas et al. “Trnp1 organizes diverse nuclear membrane-less compartments in neural stem cells”. en. In: The EMBO journal 39.16 (2020), e103373. ISSN: 0261-4189, 1460-2075. DOI: 10.15252/embj.2019103373.

[28] C. Camacho, G. Coulouris, V. Avagyan, N. Ma, J. Papadopoulos, K. Bealer, and T. L. Madden. “BLAST+: architecture and applications”. en. In: BMC bioinformatics 10.1 (2009), p. 421. ISSN: 1471-2105. DOI: 10.1186/1471-2105-10-421.

[29] A. Löytynoja. “Phylogeny-aware alignment with PRANK and PAGAN”. In: Methods in Molecular Biology. Methods in molecular biology (Clifton, N.J.) New York, NY: Springer US, 2021, pp. 17–37. DOI: 10.1007/978-1-0716-1036-7\_2.

[30] Z. Yang. “PAML: a program package for phylogenetic analysis by maximum likelihood”. en. In: Computer applications in the biosciences: CABIOS 13.5 (1997), pp. 555–556. ISSN: 0266-7061. DOI: 10.1093/bioinformatics/13.5.555.

[31] K. Zilles, E. Armstrong, K. H. Moser, A. Schleicher, and H. Stephan. “Gyrification in the cerebral cortex of primates”. en. In: Brain, behavior and evolution 34.3 (1989), pp. 143–150. ISSN: 0006-8977. DOI: 10.1159/000116500.

[32] S. H. Montgomery, N. I. Mundy, and R. A. Barton. “Brain evolution and development: adaptation, allometry and constraint”. en. In: Proc. Biol. Sci. 283.1838 (2016).

[33] T. Ohta. “Very slightly deleterious mutations and the molecular clock”. en. In: Journal of molecular evolution 26.1-2 (1987), pp. 1–6. ISSN: 0022-2844. DOI: 10.1007/BF02111276.

[34] M. Lynch and B. Walsh. The origins of genome architecture. Vol. 98. Sinauer Associates Sunderland, MA, 2007.

[35] E. Figuet, B. Nabholz, M. Bonneau, E. Mas Carrio, K. Nadachowska-Brzyska, H. Ellegren, and N. Galtier. “Life History Traits, Protein Evolution, and the Nearly Neutral Theory in Amniotes”. en. In: Molecular biology and evolution 33.6 (2016), pp. 1517–1527. ISSN: 0737-4038, 1537-1719. DOI: 10.1093/molbev/msw033.

[36] N. Lartillot and R. Poujol. “A phylogenetic model for investigating correlated evolution of substitution rates and continuous phenotypic characters”. en. In: Molecular biology and evolution 28.1 (2011), pp. 729–744. ISSN: 0737-4038, 1537-1719. DOI: 10.1093/molbev/msq244.

[37] F. Inoue and N. Ahituv. “Decoding enhancers using massively parallel reporter assays”. en. In: Genomics 106.3 (2015), pp. 159–164. ISSN: 0888-7543, 1089-8646. DOI: 10.1016/j.ygeno.2015.06.005.

[38] B. E. Bernstein et al. “The NIH Roadmap Epigenomics Mapping Consortium”. en. In: Nature biotechnology 28.10 (2010), pp. 1045–1048. ISSN: 1087-0156, 1546-1696. DOI: 10.1038/nbt1010-1045.

[39] J. Geuder, L. E. Wange, A. Janjic, J. Radmer, P. Janssen, J. W. Bagnoli, S. Müller, A. Kaul, M. Ohnuki, and W. Enard. “A non-invasive method to generate induced pluripotent stem cells from primate urine”. en. In: Scientific reports 11.1 (2021), p. 3516. ISSN: 2045-2322. DOI: 10.1038/s41598-021-82883-0.

[40] C. G. Danko et al. “Dynamic evolution of regulatory element ensembles in primate CD4+ T cells”. en. In: Nature ecology & evolution 2.3 (2018), pp. 537–548. ISSN: 2397-334X. DOI: 10.1038/s41559-017-0447-5.

[41] C. Berthelot, D. Villar, J. E. Horvath, D. T. Odom, and P. Flicek. “Complexity and conservation of regulatory landscapes underlie evolutionary resilience of mammalian gene expression”. en. In: Nature ecology & evolution 2.1 (2018), pp. 152–163. ISSN: 2397-334X. DOI: 10.1038/s41559-017-0377-2.

[42] C. D. Huber, B. Y. Kim, and K. E. Lohmueller. “Population genetic models of GERP scores suggest pervasive turnover of constrained sites across mammalian evolution”. en. In: PLoS genetics 16.5 (2020), e1008827. ISSN: 1553-7390, 1553-7404. DOI: 10.1371/journal.pgen.1008827.

[43] S. K. Reilly, J. Yin, A. E. Ayoub, D. Emera, J. Leng, J. Cotney, R. Sarro, P. Rakic, and J. P. Noonan. “Evolutionary genomics. Evolutionary changes in promoter and enhancer activity during human corticogenesis”. en. In: Science 347.6226 (2015), pp. 1155–1159. ISSN: 0036-8075, 1095-9203. DOI: 10.1126/science.1260943.

[44] M. C. Frith, M. C. Li, and Z. Weng. “Cluster-Buster: Finding dense clusters of motifs in DNA sequences”. en. In: Nucleic acids research 31.13 (2003), pp. 3666–3668. ISSN: 0305-1048, 1362-4962. DOI: 10.1093/nar/gkg540.

[45] M. Volpe, S. Shpungin, C. Barbi, G. Abrham, H. Malovani, R. Wides, and U. Nir. “trnp: A conserved mammalian gene encoding a nuclear protein that accelerates cell-cycle progression”. en. In: DNA and cell biology 25.6 (2006), pp. 331–339. ISSN: 1044-5498. DOI: 10.1089/dna.2006.25.331.

[46] S. B. Carroll. “Evo-devo and an expanding evolutionary synthesis: a genetic theory of morphological evolution”. en. In: Cell 134.1 (2008), pp. 25–36. ISSN: 0092-8674, 1097-4172. DOI: 10.1016/j.cell.2008.06.030.

[47] H. E. Hoekstra and J. A. Coyne. “The locus of evolution: evo devo and the genetics of adaptation”. en. In: Evolution; international journal of organic evolution 61.5 (2007), pp. 995–1016. ISSN: 0014-3820. DOI:10.1111/j.1558-5646.2007.00105.x.

[48] S. Tavano, E. Taverna, N. Kalebic, C. Haffner, T. Namba, A. Dahl, M. Wilsch-Bräuninger, J. T. M. L. Paridaen, and W. B. Huttner. “Insm1 Induces Neural Progenitor Delamination in Developing Neocortex via Downregulation of the Adherens Junction Belt-Specific Protein Plekha7”. en. In: Neuron 97.6 (2018), 1299–1314.e8. ISSN: 0896-6273, 1097-4199. DOI: 10.1016/j.neuron.2018.01.052.

[49] A. E. Trevino et al. “Chromatin and gene-regulatory dynamics of the developing human cerebral cortex at single-cell resolution”. en. In: Cell 184.19 (2021), 5053–5069.e23. ISSN: 0092-8674, 1097-4172. DOI: 10.1016/j.cell.2021.07.039.

[50] L. de la Torre-Ubieta, J. L. Stein, H. Won, C. K. Opland, D. Liang, D. Lu, and D. H. Geschwind. “The Dynamic Landscape of Open Chromatin during Human Cortical Neurogenesis”. en. In: Cell 172.1-2 (2018), 289–304.e18. ISSN: 0092-8674, 1097-4172. DOI: 10.1016/j.cell.2017.12.014.

[51] R. G. Arzate-Mejía, F. Recillas-Targa, and V. G. Corces. “Developing in 3D: the role of CTCF in cell differentiation”. en. In: Development 145.6 (2018), dev137729. ISSN: 0950-1991, 1477-9129. DOI: 10.1242/dev.137729.

[52] L. A. Watson, X. Wang, A. Elbert, K. D. Kernohan, N. Galjart, and N. G. Bérubé. “Dual effect of CTCF loss on neuroprogenitor differentiation and survival”. en. In: The Journal of neuroscience: the official journal of the Society for Neuroscience 34.8 (2014), pp. 2860–2870. ISSN: 0270-6474, 1529-2401. DOI: 10.1523/JNEUROSCI.3769-13.2014.

[53] D. Wu, T. Li, Z. Lu, W. Dai, M. Xu, and L. Lu. “Effect of CTCF-binding motif on regulation of PAX6 transcription”. en. In: Investigative ophthalmology & visual science 47.6 (2006), pp. 2422–2429. ISSN: 0146-0404. DOI: 10.1167/iovs.05-0536.

[54] P. Delgado-Olguín, K. Brand-Arzamendi, I. C. Scott, B. Jungblut, D. Y. Stainier, B. G. Bruneau, and F. Recillas-Targa. “CTCF promotes muscle differentiation by modulating the activity of myogenic regulatory factors”. en. In: The Journal of biological chemistry 286.14 (2011), pp. 12483–12494. ISSN: 0021-9258, 1083-351X. DOI: 10.1074/jbc.M110.164574.

[55] A. Rhie et al. “Towards complete and error-free genome assemblies of all vertebrate species”. en. In: Nature 592.7856 (2021), pp. 737–746. ISSN: 0028-0836, 1476-4687. DOI: 10.1038/s41586-021-03451-0.

[56] M. I. A. Cavassim, Z. Baker, C. Hoge, M. H. Schierup, M. Schumer, and M. Przeworski. “PRDM9 losses in vertebrates are coupled to those of paralogs *ZCWPW1* and *ZCWPW2*”. en. In: Proceedings of the National Academy of Sciences of the United States of America 119.9 (2022). ISSN: 0027-8424, 1091-6490. DOI: 10.1073/pnas.2114401119.

[57] T. Stephan et al. “Darwinian genomics and diversity in the tree of life”. en. In: Proceedings of the National Academy of Sciences of the United States of America 119.4 (2022). ISSN: 0027-8424, 1091-6490. DOI: 10.1073/pnas.2115644119.

[58] N. Jourjine and H. E. Hoekstra. “Expanding evolutionary neuroscience: insights from comparing variation in behavior”. en. In: Neuron 109.7 (2021), pp. 1084–1099. ISSN: 0896-6273, 1097-4199. DOI: 10.1016/j.neuron.2021.02.002.

[59] S. D. Smith, M. W. Pennell, C. W. Dunn, and S. V. Edwards. “Phylogenetics is the New Genetics (for Most of Biodiversity)”. en. In: Trends in ecology & evolution 35.5 (2020), pp. 415–425. ISSN: 0169-5347, 1872-8383. DOI: 10.1016/j.tree.2020.01.005.

[60] R. Chari and G. M. Church. “Beyond editing to writing large genomes”. en. In: Nature reviews. Genetics 18.12 (2017), pp. 749–760. ISSN: 1471-0056, 1471-0064. DOI: 10.1038/nrg.2017.59.

[61] W. Enard. “Functional primate genomics–leveraging the medical potential”. en. In: Journal of molecular medicine 90.5 (2012), pp. 471–480. ISSN: 0946-2716, 1432-1440. DOI: 10.1007/s00109-012-0901-4.

[62] G. Housman and Y. Gilad. “Prime time for primate functional genomics”. en. In: Current opinion in genetics & development 62 (2020), pp. 1–7. ISSN: 0959-437X, 1879-0380. DOI: 10.1016/j.gde.2020.04.007.

[63] J. Vierstra et al. “Mouse regulatory DNA landscapes reveal global principles of cis-regulatory evolution”. en. In: Science 346.6212 (2014), pp. 1007–1012. ISSN: 0036-8075, 1095-9203. DOI: 10.1126/science.1246426.

[64] F. J. Sedlazeck, P. Rescheneder, and A. von Haeseler. “NextGenMap: fast and accurate read mapping in highly polymorphic genomes”. en. In: Bioinformatics 29.21 (2013), pp. 2790–2791. ISSN: 1367-4803, 1367-4811. DOI: 10.1093/bioinformatics/btt468.

[65] S. John, P. J. Sabo, R. E. Thurman, M.-H. Sung, S. C. Biddie, T. A. Johnson, G. L. Hager, and J. A. Stamatoyannopoulos. “Chromatin accessibility pre-determines glucocorticoid receptor binding patterns”. en. In: Nature genetics 43.3 (2011), pp. 264–268. ISSN: 1061-4036, 1546-1718. DOI: 10.1038/ng.759.

[66] W. J. Kent. “BLAT–the BLAST-like alignment tool”. en. In: Genome research 12.4 (2002), pp. 656–664. ISSN: 1088-9051, 1549-5469. DOI: 10.1101/gr.229202.

[67] J. Ye, G. Coulouris, I. Zaretskaya, I. Cutcutache, S. Rozen, and T. L. Madden. “Primer-BLAST: a tool to design target-specific primers for polymerase chain reaction”. en. In: BMC bioinformatics 13 (2012), p. 134. ISSN: 1471-2105. DOI: 10.1186/1471-2105-13-134.

[68] D. A. Hysom, P. Naraghi-Arani, M. Elsheikh, A. C. Carrillo, P. L. Williams, and S. N. Gardner. “Skip the alignment: degenerate, multiplex primer and probe design using K-mer matching instead of alignments”. en. In: PloS one 7.4 (2012), e34560. ISSN: 1932-6203. DOI: 10.1371/journal.pone.0034560.

[69] G. Renaud, U. Stenzel, T. Maricic, V. Wiebe, and J. Kelso. “deML: robust demultiplexing of Illumina sequences using a likelihood-based approach”. en. In: Bioinformatics 31.5 (2015), pp. 770–772. ISSN: 1367-4803, 1367-4811. DOI: 10.1093/bioinformatics/btu719.

[70] M. Martin. “Cutadapt removes adapter sequences from high-throughput sequencing reads”. en. In: EMBnet.journal 17.1 (2011), pp. 10–12. ISSN: 2226-6089, 2226-6089. DOI: 10.14806/ej.17.1.200.

[71] M. G. Grabherr et al. “Full-length transcriptome assembly from RNA-Seq data without a reference genome”. en. In: Nature biotechnology 29.7 (2011), pp. 644–652. ISSN: 1087-0156, 1546-1696. DOI: 10.1038/nbt.1883.

[72] M. Vasimuddin, S. Misra, H. Li, and S. Aluru. “Efficient architecture-aware acceleration of BWA-MEM for multicore systems”. In: 2019 IEEE International Paralle and Distributed Processing Symposium (IPDPS). Rio de Janeiro, Brazil: IEEE, 2019, pp. 314–324. isbn: 9781728112466. DOI: 10.1109/ipdps.2019.00041.

[73] UniProt Consortium. “UniProt: a worldwide hub of protein knowledge”. en. In: Nucleic acids research 47.D1 (2019), pp. D506–D515. ISSN: 0305-1048, 1362-4962. DOI: 10.1093/nar/gky1049.

[74] O. R. P. Bininda-Emonds, M. Cardillo, K. E. Jones, R. D. E. MacPhee, R. M. D. Beck, R. Grenyer, S. A. Price, R. A. Vos, J. L. Gittleman, and A. Purvis. “The delayed rise of present-day mammals”. en. In: Nature 446.7135 (2007), pp. 507–512. ISSN: 0028-0836, 1476-4687. DOI: 10.1038/nature05634.

[75] S. Pujar et al. “Consensus coding sequence (CCDS) database: a standardized set of human and mouse protein-coding regions supported by expert curation”. en. In: Nucleic acids research 46.D1 (2018), pp. D221–D228. ISSN: 0305-1048, 1362-4962. DOI: 10.1093/nar/gkx1031.

[76] S. Henikoff and J. G. Henikoff. “Amino acid substitution matrices from protein blocks”. en. In: Proceedings of the National Academy of Sciences of the United States of America 89.22 (1992), pp. 10915–10919. ISSN: 0027-8424. DOI: 10.1073/pnas.89.22.10915.

[77] Z. Yang, R. Nielsen, N. Goldman, and A. M. Pedersen. “Codon-substitution models for heterogeneous selection pressure at amino acid sites”. en. In: Genetics 155.1 (2000), pp. 431–449. ISSN: 0016-6731. DOI: 10.1093/genetics/155.1.431.

[78] J. Felsenstein. “Phylogenies and the Comparative Method”. In: The American naturalist 125.1 (1985), pp. 1–15. ISSN: 0003-0147, 1537-5323. DOI: 10.1086/284325.

[79] E. P. Martins and T. F. Hansen. “Phylogenies and the comparative method: A general approach to incorporating phylogenetic information into the analysis of interspecific data”. In: The American naturalist 149.4 (1997), pp. 646–667. issn. 0003-0147, 1537-5323. DOI: 10.1086/286013.

[80] A. Melnikov et al. “Systematic dissection and optimization of inducible enhancers in human cells using a massively parallel reporter assay”. en. In: Nature biotechnology 30.3 (2012), pp. 271–277. ISSN: 1087-0156, 1546-1696. DOI: 10.1038/nbt.2137.

[81] F. Inoue, M. Kircher, B. Martin, G. M. Cooper, D. M. Witten, M. T. McManus, N. Ahituv, and J. Shendure. “A systematic comparison reveals substantial differences in chromosomal versus episomal encoding of enhancer activity”. en. In: Genome research 27.1 (2017), pp. 38–52. ISSN: 1088-9051, 1549-5469. DOI: 10.1101/gr.212092.116.

[82] T. Dull, R. Zufferey, M. Kelly, R. J. Mandel, M. Nguyen, D. Trono, and L. Naldini. “A third-generation lentivirus vector with a conditional packaging system”. en. In: Journal of virology 72.11 (1998), pp. 8463–8471. ISSN: 0022-538X. DOI: 10.1128/JVI.72.11.8463-8471.1998.

[83] R. Nakai, M. Ohnuki, K. Kuroki, H. Ito, H. Hirai, R. Kitajima, T. Fujimoto, M. Nakagawa, W. Enard, and M. Imamura. “Derivation of induced pluripotent stem cells in Japanese macaque (Macaca fuscata)”. en. In: Scientific reports 8.1 (2018), p. 12187. ISSN: 2045-2322. DOI: 10.1038/s41598-018-30734-w.

[84] S. Parekh, C. Ziegenhain, B. Vieth, W. Enard, and I. Hellmann. “zUMIs-A fast and flexible pipeline to process RNA sequencing data with UMIs”. en. In: GigaSciena 7.6 (2018), giy059. ISSN: 2047-217X. DOI: 10.1093/gigascience/giy059.

[85] J. W. Bagnoli, C. Ziegenhain, A. Janjic, L. E. Wange, B. Vieth, S. Parekh, J. Geuder, I. Hellmann, and W. Enard. “Sensitive and powerful single-cell RNA sequencing using mcSCRB-seq”. en. In: Nature communications 9.1 (2018), p. 2937. ISSN: 2041-1723. DOI: 10.1038/s41467-018-05347-6.

[86] A. Dobin, C. A. Davis, F. Schlesinger, J. Drenkow, C. Zaleski, S. Jha, P. Batut, M. Chaisson, and T. R. Gingeras. “STAR: ultrafast universal RNA-seq aligner”. en. In: Bioinformatics 29.1 (2013), pp. 15–21. ISSN: 1367-4803, 1367-4811. DOI: 10.1093/bioinformatics/bts635.

[87] M. I. Love, W. Huber, and S. Anders. “Moderated estimation of fold change and dispersion for RNA-seq data with DESeq2”. en. In: Genome biology 15.12 (2014), p. 550. ISSN: 1465-6906. DOI: 10.1186/s13059-014-0550-8.

[88] O. Fornes et al. “JASPAR 2020: update of the open-access database of transcription factor binding profiles”. en. In: Nucleic acids research 48.D1 (2020), pp. D87–D92. ISSN: 0305-1048, 1362-4962. DOI: 10.1093/nar/gkz1001.

[89] Alexa and Rahnenführer. “Gene set enrichment analysis with topGO”. In: Bioconductor Improv 27 (2009).

[90] R Core Team. R: A Language and Environment for Statistical Computing. Foundation for Statistical Computing. Vienna, Austria, 2019.

[91] Warnke. “Mitteilung neuer Gehirn-und Körpergewichtsbestimmungen bei Saugern”. In: Psychol. Neurol 13 (1908), pp. 355–403.

[92] E. Lewitus, I. Kelava, A. T. Kalinka, P. Tomancak, and W. B. Huttner. “An adaptive threshold in mammalian neocortical evolution”. en. In: PLoS biology 12.11 (2014), e1002000. ISSN: 1544-9173, 1545-7885. DOI: 10.1371/journal.pbio.1002000.

[93] G. Crile and D. P. Quiring. “A record of the body weight and certain organ and gland weights of 3690 animals”. In: (1940).

[94] R. T. Bronson. “Brain weight-body weight relationships in 12 species of nonhuman primates”. en. In: American journal of physical anthropology 56.1 (1981), pp. 77–81. ISSN: 0002-9483, 1096-8644. DOI: 10.1002/ajpa.1330560109.

[95] J. K. Rilling and T. R. Insel. “The primate neocortex in comparative perspective using magnetic resonance imaging”. en. In: Journal of human evolution 37.2 (1999), pp. 191–223. ISSN: 0047-2484. DOI: 10.1006/jhev.1999.0313.

[96] A. Hrdlička. “Weight of the brain and of the internal organs in American monkeys. With data on brain weight in other apes”. In: American journal of physical anthropology 8.2 (1925), pp. 201–211. ISSN: 0002-9483, 1096-8644. DOI: 10.1002/ajpa.1330080207.

[97] P. R. Manger. “An examination of cetacean brain structure with a novel hypothesis correlating thermogenesis to the evolution of a big brain”. en. In: Biological reviews of the Cambridge Philosophical Society 81.2 (2006), pp. 293–338. ISSN: 1464-7931, 0006-3231. DOI: 10.1017/S1464793106007019.

[98] K. Kverková, T. Bělíková, S. Olkowicz, Z. Pavelková, M. J. O’Riain, R. Sumbera, H. Burda, N. C. Bennett, and P. Němec. “Sociality does not drive the evolution of large brains in eusocial African mole-rats”. en. In: Scientific reports 8.1 (2018), p. 9203. ISSN: 2045-2322. DOI: 10.1038/s41598-018-26062-8.

[99] L. Ventura-Antunes, B. Mota, and S. Herculano-Houzel. “Different scaling of white matter volume, cortical connectivity, and gyrification across rodent and primate brains”. en. In: Frontiers in neuroanatomy 7 (2013), p. 3. ISSN: 1662-5129. DOI: 10.3389/fnana.2013.00003.

[100] E. A. Spitzka. “Brain-weights of animals with special reference to the weight of the brain in the Macaque monkey”. en. In: The Journal of comparative neurology 13.1 (1903), pp. 9–17. ISSN: 0021-9967. DOI: 10.1002/cne.910130103.

[101] Brodmann. “Neuere Forschungsergebnisse der Großhirnrindenanatomie mit besonderer Berücksichtigung anthropologischer Fragen”. In: The Science of Nature 1.46 (1913), pp. 1120–1122. ISSN: 0028-1042, 1432-1904.

[102] H. Stephan, H. Frahm, and G. Baron. “New and revised data on volumes of brain structures in insectivores and primates”. en. In: Folia primatologica; international journal of primatology 35.1 (1981), pp. 1–29. ISSN: 0015-5713. DOI: 10.1159/000155963.

